# Imaging of existing and newly translated proteins elucidates mechanisms of sarcomere turnover

**DOI:** 10.1101/2023.08.31.555653

**Authors:** Guy Douvdevany, Itai Erlich, Lilac Haimovich-Caspi, Tomer Mashiah, Maksymilian Prondzynski, Maria Rosaria Pricolo, Jorge Alegre-Cebollada, Wolfgang A. Linke, Lucie Carrier, Izhak Kehat

## Abstract

**Background:** How the sarcomeric complex is continuously turned-over in long-living cardiomyocytes is unclear. According to the prevailing model of sarcomere maintenance, sarcomeres are maintained by cytoplasmic soluble protein pools with free recycling between pools and sarcomeres.

**Methods:** We imaged and quantified the turnover of expressed and endogenous sarcomeric proteins, including the giant protein titin, in cardiomyocytes in culture and in vivo, at the single cell and at the single sarcomere level using pulse-chase labeling of Halo-tagged proteins with covalent ligands.

**Results:** We disprove the prevailing ‘protein pool’ model and instead show an ordered mechanism in which only newly translated proteins enter the sarcomeric complex while older ones are removed and degraded. We also show that degradation is independent of protein age, and that proteolytic extraction is a rate limiting step in the turnover. We show that replacement of sarcomeric proteins occurs at a similar rate within cells and across the heart and is slower in adult cells.

**Conclusions:** Our findings establish a ‘unidirectional replacement’ model for cardiac sarcomeres subunit replacement and identify their turnover principles.

## Introduction

Almost all biological processes depend on the assembly of proteins into functional complexes. The protein subunits of these functional complexes must be continuously replaced since mammalian cells often live much longer than their proteins.^1,2^ Two models for the assembly of complexes were previously proposed. In the first model the complex is in a dynamic equilibrium with a pool of cytoplasmic proteins and continuously recycles and exchanges subunits with the pool through de-and re-assembly.^3^ This exchange was shown to maintain simple complexes such as actin fibers and microtubules.^4,5^ An alternative model proposes a sequential assembly^6^, where the complex is formed following an ordered series of steps. For example, the 26S proteasome shows a sequential assembly.^7^ However, the mechanism by which such complexes are continuously maintained once they are fully assembled, and the role of subunit recycling in their maintenance remains largely unknown.^8^

The sarcomere is the basic contractile protein complex of striated muscle, including cardiomyocytes. It is composed of the contractile myofilament proteins myosin and actin linked by titin (TTN), along with several other regulatory, structural, and accessory proteins.^9^ Cardiomyocytes must continuously maintain their sarcomeres since sarcomeric proteins have a short half-life^10^, and the decreased sarcomere and sarcomeric protein content in failing hearts suggest that failure to do so may lead to heart failure.^11,12^ According to the prevailing model of sarcomere maintenance, sarcomeres are maintained by cytoplasmic soluble protein pools with free recycling between pools and sarcomeres.^13–15^ This model was more recently referred to as the “sarcostat”.^14,16–18^ Support for this model came from fluorescence recovery after photobleaching (FRAP) experiments with fluorescently tagged sarcomeric proteins, that show recovery of fluorescent-actin^19,20^, myotilin^21,22^, α-actinin 2 (ACTN2), telethonin, tropomyosin 1 (TPM1)^23^, and TTN^24^ after bleaching within several hours. In the latter study it was even proposed that TTN molecules can move between adjacent sarcomeric complexes.^24^

Here we developed a pulse-chase approach based on covalent binding of different color fluorescent HaloTag ligands to image the replacement of older sarcomeric proteins by newly translated ones in cardiomyocytes in culture and in vivo. We imaged and measured the turnover of overexpressed and endogenous sarcomeric proteins, including the largest known protein TTN within the sarcomeric complex, in cultured cells and in vivo, at the single cell and the single sarcomere level. In contrast to the current ‘protein pool’ model of sarcomere maintenance, our findings provide evidence for an ordered process in which newly translated proteins are continuously added to the complex, while assembled proteins are continuously removed, degraded, and not recycled. We further show that once assembled, newly synthesized proteins are just as likely to be removed and degraded as older ones, and that proteolytic cleavage is needed for their extraction from the sarcomere. Our study thus establishes the ‘unidirectional replacement’ model for cardiac sarcomeres.

## Methods

### Data Availability

The data that support the findings of this study are available from the corresponding author upon reasonable request. The detailed methods section is available in the Supplemental Material.

### Animal experiments

All animal procedures were approved by the Technion-Israel Institute of Technology Institutional Animal Care and Use Committee and conducted in accordance with guidelines for the care and use of laboratory animals. All results are reported in compliance with the Animal Research Reporting of In Vivo Experiments (ARRIVE) guidelines. Mice were randomly assigned to each experimental group.

## Results

### Sarcomere maintenance is a unidirectional process

To image protein turnover in single living cardiomyocytes, we initially expressed Halotagged sarcomeric proteins in cultured neonatal rat ventricular cardiomyocytes (NRVMs) using adenoviral vectors. For the Z-line ACTN2 and the A-band myosin regulatory light chain 2 (MYL2) sarcomeric proteins we used full-length sequence (ACTN2-Halo and Halo-MYL2), and for the large M-line myomesin-1 (MYOM1) and the A-band cardiac myosin-binding protein C (MYBPC3) we used the N and C-terminal fragments, respectively (MYOM1-Halo and Halo-MYBPC3). These fragments are small enough to be packaged in adenoviral vectors and sufficient for stable integration into sarcomeres.^25,26^ After 48 hours, to allow expression of the Halo-tagged proteins, we pulsed the cells with the Oregon Green (OG) HaloTag-ligand. HaloTag is a modified haloalkane dehalogenase that covalently and irreversibly binds to HaloTag-ligand.^27^ Therefore, once bound by the ligand the existing Halo-tagged proteins in the cell remain permanently tagged with fluorescent green. To ensure that all the Halo-tagged proteins in the cell are bound by a ligand, we followed by pulsing with the non-fluorescent blocker ligand 7-bromoheptanol (7BRO) at a high concentration (65µM). 7BRO is an irreversible color-less haloalkane with similar structure to other Halo-ligands.^28^ We then extensively washed the cells from excess ligands, photobleached a small area in the sarcomere, added the far-red HaloTag ligand Janelia Fluor 635 (JF-635)^29^, and imaged the cardiomyocytes for at least 72 hours (Figure 1A). We limited the frequency of imaging to 12-24 hours to reduce phototoxicity. The JF-635 ligand shows almost no light absorption in aqueous solution, but upon binding to HaloTag, it increases its absorbance and fluorescence by more than 100-fold, and therefore can be used for continuous live cell imaging.^30^ As expected, we did not observe far-red fluorescence signal immediately after the addition of the JF-635 ligand, confirming that all the Halo-tagged proteins in the cell were already bound, and that unbound JF-635 does not significantly fluoresce. However, once new Halo-tagged sarcomeric proteins were translated their available HaloTag could bind the JF-635 ligand and display far-red fluorescence. Our approach therefore allowed us to follow two main populations of sarcomeric proteins in the myocytes: older proteins labeled in green and newly translated sarcomeric proteins labeled in far-red. Over time, we observed an increase in the sarcomeric far-red fluorescence signal across the myocyte’s sarcomeres along with a simultaneous decrease in green fluorescence, showing that newly synthesized proteins were added to the sarcomere and gradually replaced the older ones (Figure 1B). We did not observe old or new Halo-tagged proteins outside the sarcomere, indicating that un-incorporated sarcomeric proteins are degraded.

**Figure 1.**
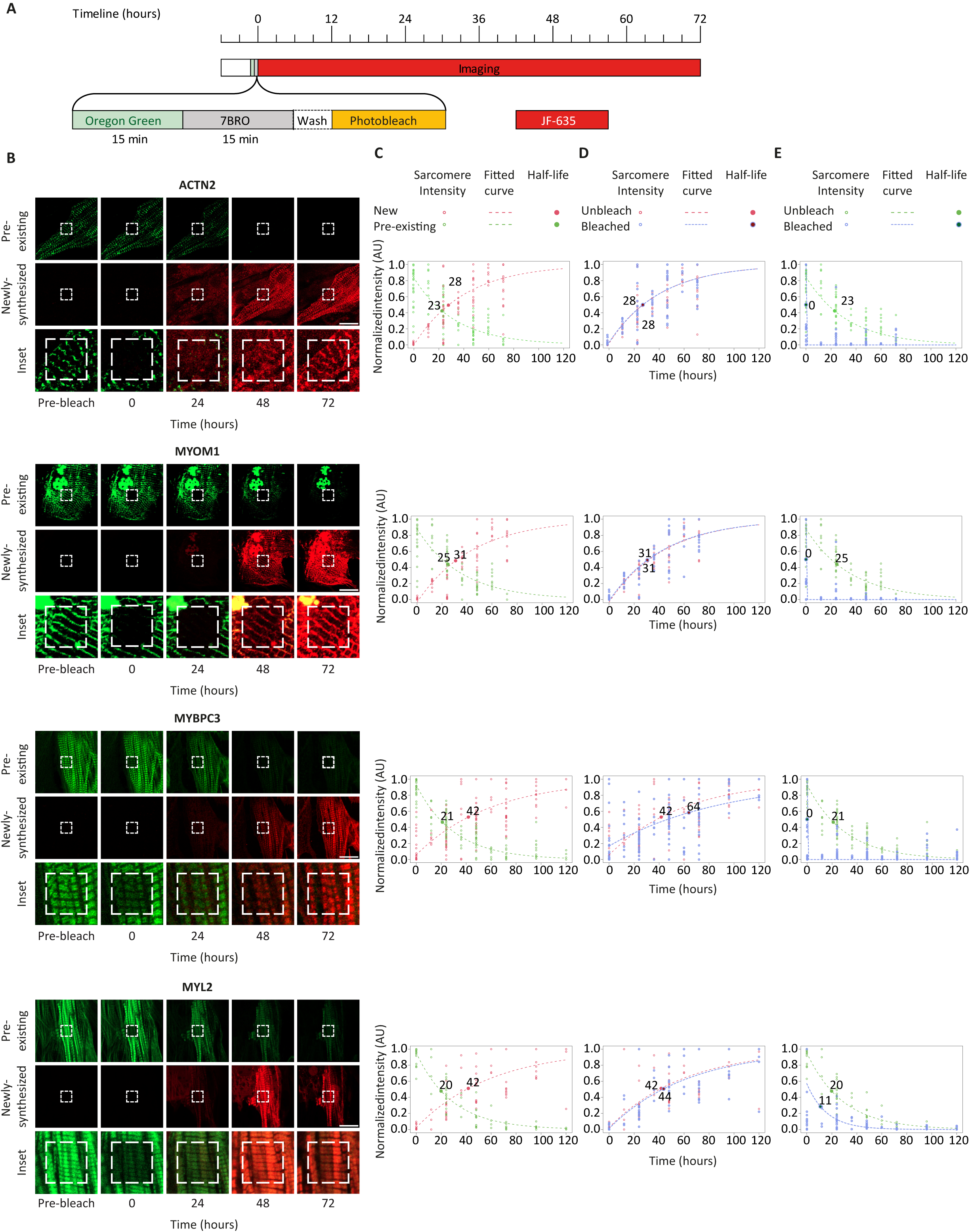
Imaging of sarcomere turnover shows a unidirectional process. **A,** Pulse-chase experiment timeline: Cardiomyocytes expressing Halo-tagged sarcomeric proteins were incubated with Oregon Green ligand, followed by a non-fluorescent blocker ligand 7-bromoheptanol (7BRO), washed, and incubated with far-red Janelia Fluor-635 HaloTag ligand (JF-635). A small square part of the cell was photobleached, followed by at least 72 hours of time-lapsed imaging. **B,** Representative time-lapse images of cardiomyocytes expressing Halo-tagged ACTN2, MYOM1, MYBPC3 and MYL2 in the green (top), far-red (middle), and the magnified merged channel (bottom), before the photobleaching, immediately after (0), and at 24,48, and 72 hours, showing gradual replacement of pre-existing proteins (green) with newly synthesized ones (far-red). The dashed white square is the photobleached field. It is gradually filling with new far-red labeled proteins but not with green labeled ones. Scale bar = 30µm. **C**, Normalized sarcomeric fluorescence amplitude is shown for the green and far-red channels with fitted exponential decay and accumulation curves (dotted lines) and calculated half-lives (bold dot and number; n=11-26 cells for each protein from N=3 independent experiments). **D**, Normalized sarcomeric fluorescence intensity is shown in the far-red channel for unbleached (light red) and bleached (light blue) myofibrils, showing similar rates of accumulation. **E**, Normalized sarcomeric fluorescence intensity is shown in the green channel showing exponential decay in the unbleached myofibril (light green) and no recovery in the bleached myofibril (light blue), except for some recovery in MYL2.

We wondered whether subunit recycling is part of the maintenance process, and therefore we photobleached a small part of the green-labeled sarcomere at the beginning of the time lapse (Figure 1A and 1B). According to the ‘protein pool’ model, sarcomeres would be maintained by a pool of cytoplasmic proteins containing a mixture of both older recycled and newly translated sarcomeric proteins. As a result, the photobleached sarcomeric proteins should be gradually replaced by both ‘old’ green labeled proteins and ‘new’ far-red labeled proteins from the pool (Figure S1). Contrary to this prediction, we found that the photobleached area gradually filled with ‘new’ far-red labeled proteins, whereas the green signal failed to recover, showing sarcomeric proteins are not recycled (Figure 1B). We initially used strong laser power to completely bleach the green signal. The green signal did not recover under these conditions, but the emergence of red florescence was somewhat retarded inside the bleached area, indicative of some local photodamage. We therefore opted for a lower level of bleaching. There was still no recovery of the green signal under these circumstances, and uniform emergence of red fluorescence was observed. To test whether we could identify diffusion we also expressed ACTN2-Halo and performed FRAP in rat fibroblasts. Green-labeled ACTN2-Halo fluorescence recovered in fibroblasts within a minute of bleaching (Figure S2), in contrast to cardiomyocytes where no recovery occurred.

To quantify our findings and measure the intensity of the green and far-red signals, we stacked the time-lapse images, drew a linear region of interest along the myofibril and calculated the median amplitude of the signal intensity over time in these two channels along the line (Figure S3). In addition to the sarcomeric signal we also observed a nuclear accumulation of signal for MYOM1, which likely represents an aggregate, and were therefore careful not to include the nucleus in our analyses. Assuming a first order kinetic^31,32^, we fitted an exponential decay function to the degradation, or a function that gives the difference between 1 and the exponential decay to fit to synthesis and calculated the sarcomeric half-lives of proteins from the fitted curves (Figure 1C). These data show that the sarcomeric half-lives calculated for ACTN2, MYOM1, MYBPC3, and MYL2 were similar. Our approach does not require the assumption of a steady state for quantification, and we could calculate the half-life from either the degradation rate of green labeled proteins or from the accumulation of far-red tagged proteins. While these calculations yielded values in the same range, the half-lives calculated from accumulation were longer than those calculated from degradation. Some correction factor is likely to be required, and bleaching from the timelapse imaging could have contributed to accelerating green fluorescence decline and delaying far-red fluorescence accumulation. Together these analyses show that proteins are turned over in the sarcomere with a half-life of 20-42 hours in cultured cardiomyocytes.

We measured and compared the increase in incorporation of newly translated far-red labeled proteins outside and inside the bleached area and found similar rates, showing that sarcomeric proteins in different myofibrils are replaced simultaneously and at similar rates, and that our photobleaching did not damage the sarcomere (Figure 1D). Similarly, we measured and compared the changes in the existing, green-labeled protein signal outside and inside the bleached area (Figure 1E). All the protein studied showed exponential decline in intensity outside the bleached area and for ACTN2, MYOM1 and MYBPC3 we did not observe any recovery of the green signal in the bleached area. The exception was a small resurgence of green fluorescence immediately after photobleaching for MYL2 followed by an exponential decay (Figure 1E), which may represent a small mobile unincorporated pool of MYL2. However, even if such a MYL2 pool existed it would be relatively small. Clearly, most of the existing MYL2 proteins in the sarcomere were replaced by newly synthesized ones and not through recycling. We could also quantify the turnover of single sarcomeres using super-resolution. To this end we tracked several single sarcomeres in individual cells and measured their signal amplitude in the stacked time lapse images. We fitted an exponential decay function to the measured data and calculated the half-life for each sarcomere individually. Although some variability existed, the half-lives of individual sarcomeres within the same cell were similar (Figure S4). To further validate our half-life measurements, we transduced NRVMs with ACTN2-Halo using adenoviral vector. We similarly pulsed the cells with OG HaloTag-ligand followed by pulsing with 7BRO. We then extensively washed thecells from excess ligands and waited 0,24, or 48 hours. At the end of the waiting time, we pulsed the cells with red TMRDirect Halo-Ligand and harvested the cultures. The NRVM protein extract was analyzed using SDS-PAGE with direct measurements of fluorescence from the gel. This analysis showed similar half-lives for ACTN2 (Figure S5A through S5C). The half-life of ACTN2 was also estimated following blockade of translation using cycloheximide for 8 hours in NRVMS, showing a similar half-life (Figure S5D and S5E).

Collectively, our data refute the prevailing ‘protein pool’ hypothesis and prove that sarcomere maintenance is unidirectional: new proteins are incorporated into the sarcomere, but older proteins are extruded and degraded and cannot be recycled. Our measurements of the half-lives also imply that sarcomeric proteins are replaced with similar rates throughout the myocyte sarcomeres.

### Imaging turnover of endogenous sarcomeric proteins

To confirm that our turn-over model also applies to endogenous cardiac proteins we used neonatal mice ventricular cardiomyocytes (NMVMs) derived from animals in which HaloTag was knocked-in to the I-band section of the largest known protein, the sarcomeric protein TTN.^33^ We used the same protocol as used in NRVMs to image the turnover of endogenous TTN in mouse cardiomyocytes. Similar to NRVMs we used minimal sufficient bleaching in our experiments because we found that harsh bleaching damaged the cells. As we showed for over-expressed sarcomeric proteins, the pre-existing TTN proteins were gradually and uniformly replaced by newly translated ones across the cardiomyocyte (Figure 2A). After photobleaching there was no reemergence of the green fluorescence showing that endogenous TTN cannot be recycled (Figure 2A). Quantification of our data confirmed this observation and showed that TTN is turned over with a half-life of about 15-31 hrs. (Figure 2B through 2D).

**Figure 2.**
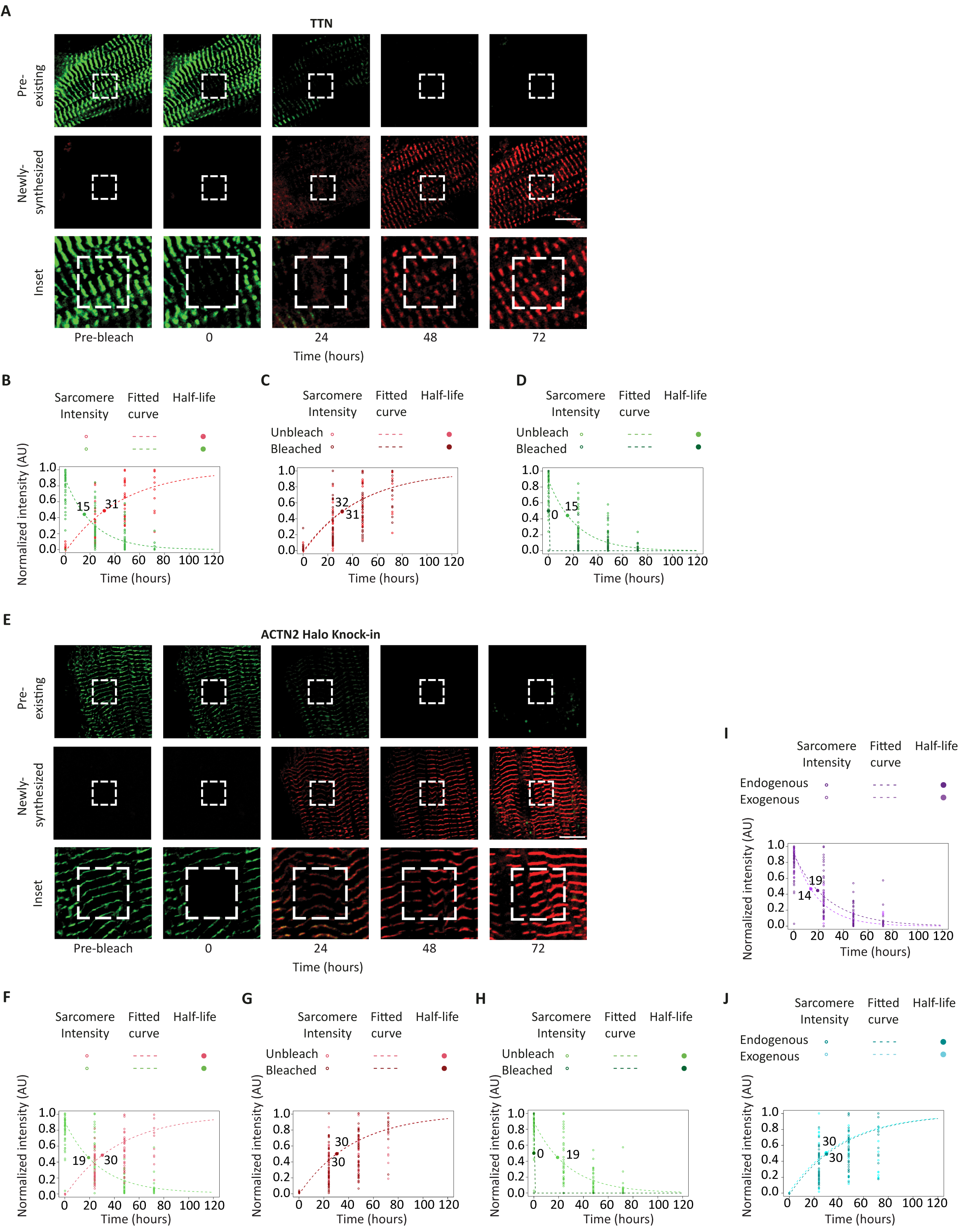
Imaging turnover of the endogenous sarcomeric proteins TTN and ACTN2. **A,** Representative time-lapse images of neonatal mouse ventricular cardiomyocytes from Halo-tagged TTN knock-in mice in the green (top), far-red (middle), and in magnified merged inset (bottom), before the photobleaching, immediately after (0), and at 24,48, and 72 hours, showing gradual replacement of pre-existing TTN proteins (green) with newly synthesized ones (far-red). The dashed white square is the photobleached field. It is gradually filling with new far-red labeled proteins but not with green labeled ones. Scale bar = 30µm. **B**, Normalized sarcomeric fluorescence amplitude is shown for the green and far-red channels with fitted exponential decay and accumulation curves (dotted lines) and calculated half-lives (bold dot and number). n=41 cells from N=3 independent experiments. **C**, Normalized sarcomeric fluorescence intensity is shown in the far-red channel for unbleached (light red) and bleached (dark red) myofibrils, showing similar degree of accumulation. **D**, Normalized sarcomeric fluorescence intensity is shown in the green channel showing exponential decay in the unbleached myofibril (light green) and no recovery in the bleached myofibril (dark green). **E**, Representative time-lapse images of NMVM after HITI knock-in of HaloTag to the *Actn2* locus using the same protocol as in (A), showing gradual replacement of existing endogenous ACTN2 proteins (green) with newly translated ones (far-red). The photobleached field is gradually filling with new far-red labeled proteins but not with green labeled ones. **F-H**, Quantification and calculation of assembly and degradation half-lives for endogenous ACTN2 as in (**B-D**). n=44 cells from N=3 independent experiments. **I-J**, Comparison of turnover rates of endogenous knocked-in ACTN2-Halo to exogenous virally expressed ACTN2-Halo in NMVMs. Normalized sarcomeric fluorescence amplitude is shown for the green (**I**) and far-red (**J**) channels with fitted exponential decay and accumulation curves (dotted lines) and calculated half-lives (bold dot and number). The deep and light purple dots and lines in (**I**) show the degradation of endogenous and exogenous virally expressed ACTN2 from the green channel respectively. The deep and light cyan dots and lines in (**J**) show the accumulation of endogenous and exogenous virally expressed ACTN2 from the far-red channel. n=44 endogenous HITI cardiomyocytes and n=22 exogenous virally transduced cardiomyocytes, each from N=3 independent experiments.

Next, we used an approach that allowed us to knock in the HaloTag to endogenous genes in cultured cardiomyocytes. The CRISPR/Cas9-based homology-independent targeted integration (HITI) strategy can generate precise and efficient knock-in in genes of non-dividing cells like cardiomyocytes by using non-homologous end-joining, with no insertions or deletions at most junction sites.^34^ To knock in the HaloTag in the endogenous *Actn2* locus in cardiomyocytes using HITI we identified a unique Cas9 target site upstream of the C-terminal stop codon of the *Actn2* gene. We used the same Cas9 target site to flank the donor integration cassette, which contained the *Actn2* C-terminus without a stop codon, followed in-frame by the HaloTag, allowing the same single guide RNA (sgRNA) to guide Cas9 to cut both in the genomic and on both sides of the cassette (Figure S6A). Since integration of the cassette would regenerate the sgRNA site and allow Cas9 to cut the cassette out again, the Cas9 cut sites flanking the cassette were placed inversely to the sgRNA orientation in the genome, creating a hybrid junction that the sgRNA can no longer recognize after integration. We isolated NMVMs from Cas9-expressing mice^35^, transduced them with adeno associated viruses (AAVs) encoding for the donor cassette and for either the *Actn2* C-terminal sgRNA or for control non-targeting sgRNA. Strong Z-line staining was observed after the HaloTag ligand addition in ∼15% of cardiomyocytes only when *Actn2* sgRNA was used, and not when control non-targeting sgRNA was used, indicating that HaloTag cassette was integrated properly in frame in these cardiomyocytes (Figure S6B and S6C). SDS-PAGE analysis confirmed the expression of ACTN2-Halo at the appropriate molecular weight albeit with lower levels compared to adeno-virally over-expressed ACTN2-Halo (Figure S6D and S6E). We executed the same imaging protocol we used for TTN and for expressed Halo tagged proteins and followed the turnover of endogenous ACTN2 with knocked-in HaloTag. The turnover of endogenous ACTN2 in NMVMs was similar to what we observed with virally expressed ACTN2-Halo in rat cardiomyocytes: pre-existing ACTN2 proteins were gradually and uniformly replaced by newly translated ones across the cardiomyocyte, and after photobleaching there was no reemergence of the green fluorescence, showing that endogenous ACTN2 cannot be recycled (Figure 2E). Quantification of the data showed that endogenous ACTN2 is turned over with a half-life of about 19-31 hrs. (Figure 2F through 2H). Together our data on ACTN2 and TTN demonstrated that both overexpressed and endogenous proteins cannot be recycled.

We also transduced NMVMs with adenoviral vectors encoding for ACTN2-Halo and compared the turnover rates of exogenously over-expressed ACTN2-Halo to endogenous HITI generated knock-in ACTN2-Halo (Figure 2I and 2J). We found that these turnover rates were similar, implying that expression level does not determine the turnover of sarcomeric proteins in the complex. We used the same HITI approach to knock-in Halo to the C-terminus of endogenous *Myom1* and calculated the half-life, showing similar value to the one calculated with over-expressed N-terminal fragment of MYOM1 (Figure S6F and S6G). Here we did not observe a nuclear accumulation suggesting it resulted from either the expression of an N-terminal fragment or from the higher expression level achieved using adenoviral vectors.

### Sarcomere protein degradation is independent of protein age and controls turnover rate

Protein degradation is mainly thought to be stochastic, although it has been hypothesized that old sarcomeric proteins are identified by quality control pathways and targeted for degradation.^31,32,36^ We therefore wanted to determine whether young and old proteins in the sarcomere are turned over at the same rate. We used MYOM1-Halo that has a very strong signal and expressed it in NRVM using an adenoviral vector. After 48 hours, we incubated the cells with the OG ligand and then with the JF-635 ligand for an additional 48 hours. To block all HaloTags we pulsed with the non-fluorescent blocker 7BRO at a high concentration (65µM) for 15 minutes after each fluorescent ligand, followed by extensive washing. This labeling protocol allowed us to follow the degradation of two main populations of sarcomeric proteins in the myocytes: older proteins labeled green and newer proteins labeled far-red over the next 72 hours (Figure 3A). As before, we were careful not to include the MYOM1 nuclear aggregate in our analyses. We observed a gradual decline in the sarcomeric signal intensity for both the old and the new proteins (Figure 3B). We fitted the signal data to an exponential decay function (Figure S7A) and found similar rates of decay and a similar half-life for both the old and new MYOM1 proteins (Figure 3C and 3D). We also used an alternative protocol in which cells were similarly labeled with OG ligand 48 hours after the transduction, but were then incubated with 7BRO for 12 hours, followed by washing and labeling with JF-635 ligand for 24 hours, ensuring that the youngest green labeled proteins are at least 12 hours older than the oldest red labeled ones (Figure S7B). Following these cells for 72 hours showed that the half-lives of the old and new MYOM1 proteins were similar as well (Figure S7C through S7E). These results support a stochastic model where all proteins have the same likelihood of being removed and degraded, regardless of how old they are.

**Figure 3.**
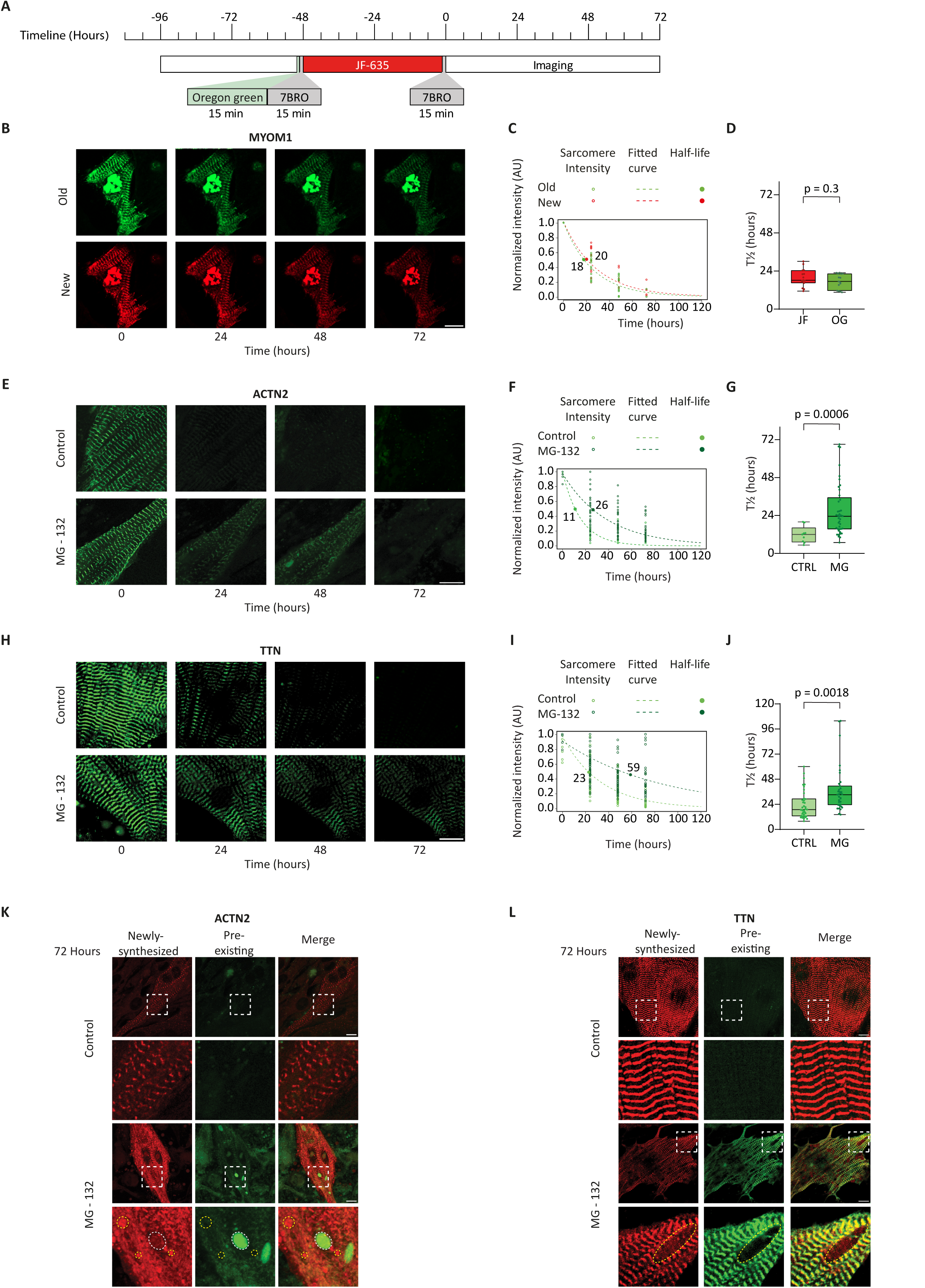
Degradation is independent of protein age and requires proteases. **A,** Degradation experiment timeline - Cardiomyocytes expressing MYOM1-Halo for 48 hours were incubated with the OG and then with the JF-635 Halo-ligands for an additional 48 hours and imaged in time-lapse 72 hours. **B,** Representative images of cardiomyocytes at 0,24, 48, and 72 hours show a gradual decline of the sarcomeric signal in both the green and far-red channels. Scale bar = 10.25µm. **C-D,** Quantification of normalized sarcomeric fluorescence intensity in the green (OG) and far-red (JF) channels (**C**), showing the half-lives of newer and older proteins in individual cardiomyocytes did not differ (**D**) (n=15 cells, N=2 independent experiments, two tailed paired Student’s t-test p=0.3). **E**, Representative time-lapse images of NRVMs expressing ACTN2-Halo incubated with (bottom) or without (top) MG-132 at 0, 24, 48, and 72 hours in the green channel show that inhibition of proteases by MG-132 resulted in a marked delay in the removal of existing OG labeled proteins from the sarcomere. Scale bar=10µm **F-G**, Quantification of normalized sarcomeric fluorescence intensity for control (light green, Ctrl) and MG-132 treated cardiomyocytes (dark green) (**F**) show much longer half-lives of ACTN2-Halo in MG-132-treated than in control cardiomyocytes (**G**) (n=9-41 cells, N=3 independent experiments, two tailed Student’s t-test p<0.001). **H-J** similar analyses as in (**E-G**) in NMVMs from Halo-TTN knock-in mice show much longer half-lives of endogenous Halo-TTN in MG-132-treated cardiomyocytes relative to controls. Scale bar = 10µm (n=37-35 cells, N=3 independent experiments, two tailed Student’s t-test p<0.01). **K**, Representative images of cardiomyocytes expressing ACTN2-Halo treated with control or MG-132 after 72 hours in the merged green and far-red channel with inset (dashed square) showing accumulation of newly translated far-red labeled proteins outside the Z-line and multiple aggregates of newly translated (yellow dotted circle) proteins and some aggregates of old proteins (white dotted circle). Scale bar = 10µm. **L**, Representative images of cardiomyocytes with knock-in Halo-TTN treated with control or MG-132 after 72 hours in the merged green and far-red channels with inset (dashed square) showing aggregates composed of newly translated far-red labeled proteins (yellow dotted circle) that displace the myofibrils. Scale bar = 10µm.

Both calpain and proteasomal proteases are responsible for the degradation of sarcomeric proteins.^36^ However, it is not known if the proteolytic activity of these enzymes is required for the removal of proteins from the sarcomeric complex, or whether sarcomeric proteins can detach intact from the complex and are only degraded later. To differentiate between these two hypotheses, we used the peptide aldehyde MG-132, which is capable of inhibiting both calpains and the proteasome.^37^ We expressed ACTN2-Halo in NRVMs using adenoviral vector and repeated the imaging protocol we used to study the turnover and measured the half-life of ACTN2-Halo with or without incubation with MG-132. The inhibition of proteases by MG-132 resulted in a marked delay in the removal of existing green-labeled ACTN2-Halo proteins from the sarcomeric structure with a significant increase in ACTN2 sarcomeric half-life (Figure 3E through 3G). We validated the effect of MG-132 using a Western blot, showing an increase in ubiquitinated proteins (Figure S8A), and used it to estimate the half-life of endogenous ACTN2, further confirming our imaging analysis (Figure S8B and S8C). We performed the same imaging analysis with endogenous TTN in NMVMs of the Halo-TTN knock-in mice, and similarly observed a marked delay in the removal of existing TTN and a significant increase in TTN sarcomeric half-life (Figure 3H through 3J). To specifically assess the role of calpains we used the calpain-1 inhibitor N-acetyl-leucyl-leucyl-norleucinal (ALLN)^38^ instead of MG-132. We show that like with MG-132, the inhibition by ALLN results in a significant delay in the removal of endogenous Halo-TTN from the sarcomere with increased half-life (Figures S8D through S8F). This delayed turnover indicates that proteolytic activity is required for the removal of sarcomeric proteins such as ACTN2 and TTN from the sarcomeric structure.

Earlier, we posited that sarcomeric proteins are translated abundantly, accompanied by swift proteasomal degradation of any unincorporated proteins.^39^ In agreement with this proposal, we found that newly translated ACTN2-Halo proteins accumulated outside the Z-line structure following proteolytic blockade. By 72 hours multiple aggregates composed predominantly of newly translated far-red-labeled proteins were observed (Figure 3K and Figure S9A). Interestingly we also detected several discreet large aggregates predominantly made of older green labeled ACTN2-Halo proteins (Figure 3K). These aggregates could contain proteins derived from the disintegrating sarcomeric structure observed at this stage. Unlike ACTN2-Halo which was over-expressed in these experiments, the Halo-TTN is an endogenous protein. Large aggregates composed of newly translated far-red labeled Halo-TTN were observed between the myofibrils after proteolytic blockade, but these accumulations were not as dense as those of ACTN2-Halo (Figure 3I and Figure S9B). Together these data show that when proteolysis is blocked the removal of sarcomeric proteins such as ACTN2 and TTN from the sarcomere is delayed and the half-life of these proteins in the structure is increased. Newly translated proteins cannot push the existing intact proteins out of the structure without proteolysis, and if newly translated proteins cannot be integrated or degraded, they accumulate outside the sarcomere. These findings suggest that proteolysis is the rate limiting step in sarcomere turnover.

### Imaging protein turnover in vivo using viral vectors

To image and quantify the turnover of sarcomeric proteins in vivo in the adult mouse heart we transduced neonatal mice with AAV vectors encoding TPM1-Halo or ACTN2-Halo full length proteins. After waiting for the mice to reach the age of 4 months we injected the non-fluorescent blocker 7BRO intravenously and intraperitoneally to ensure saturation of all the available HaloTags. We then waited either 3 hours, 3, or 7 days and intravenously delivered the fluorescent Janelia Fluor-525 HaloTag ligand (JF-525). For TPM1-Halo we also included a 1-day group. To measure the maximal possible labeling intensity, we also delivered Halo-ligand JF-525 to mice that did not receive the 7BRO blocker. In addition to the ‘3 hours’ group we included another negative control group of mice that received the ligands but not the AAVs encoding for Halo-tagged proteins (Figure 4A). We harvested the hearts 40 minutes after the injection of the JF-525 Halo-ligand and sectioned. Low and high magnification images of cardiac sections showed background signal in the ‘3 hours’ negative control group (Figure 4B through 4E); the sarcomeric fluorescence signal was increased by 3 days, and more so after 7 days, and was highest in the ‘no 7BRO’ cardiomyocytes. In the latter, it was evident that the transduction rates are significant, albeit with some cell-to-cell variability. As we did in the cultured cardiomyocytes, we drew a region of interest along the myofibril in individual cardiomyocytes in the sections (Figure 4D through 4G) and calculated the median amplitude of the sarcomeric signal (Figure 4H and 4I). We fitted the data to a 1 minus exponential decay function and estimated the half-life of ACTN2 and TPM1 to be 285 and 108 hours, respectively (Figure 4J and 4K). Although we could not follow the same cardiomyocytes over time in vivo as we could in culture experiments, we could estimate the half-life from the average of multiple cells. We found that the measured half-life of ACTN2 in vivo was ∼10-fold longer than in cultured neonatal cardiomyocytes, indicating a slowing of the turn-over as cardiomyocytes matured.

**Figure 4.**
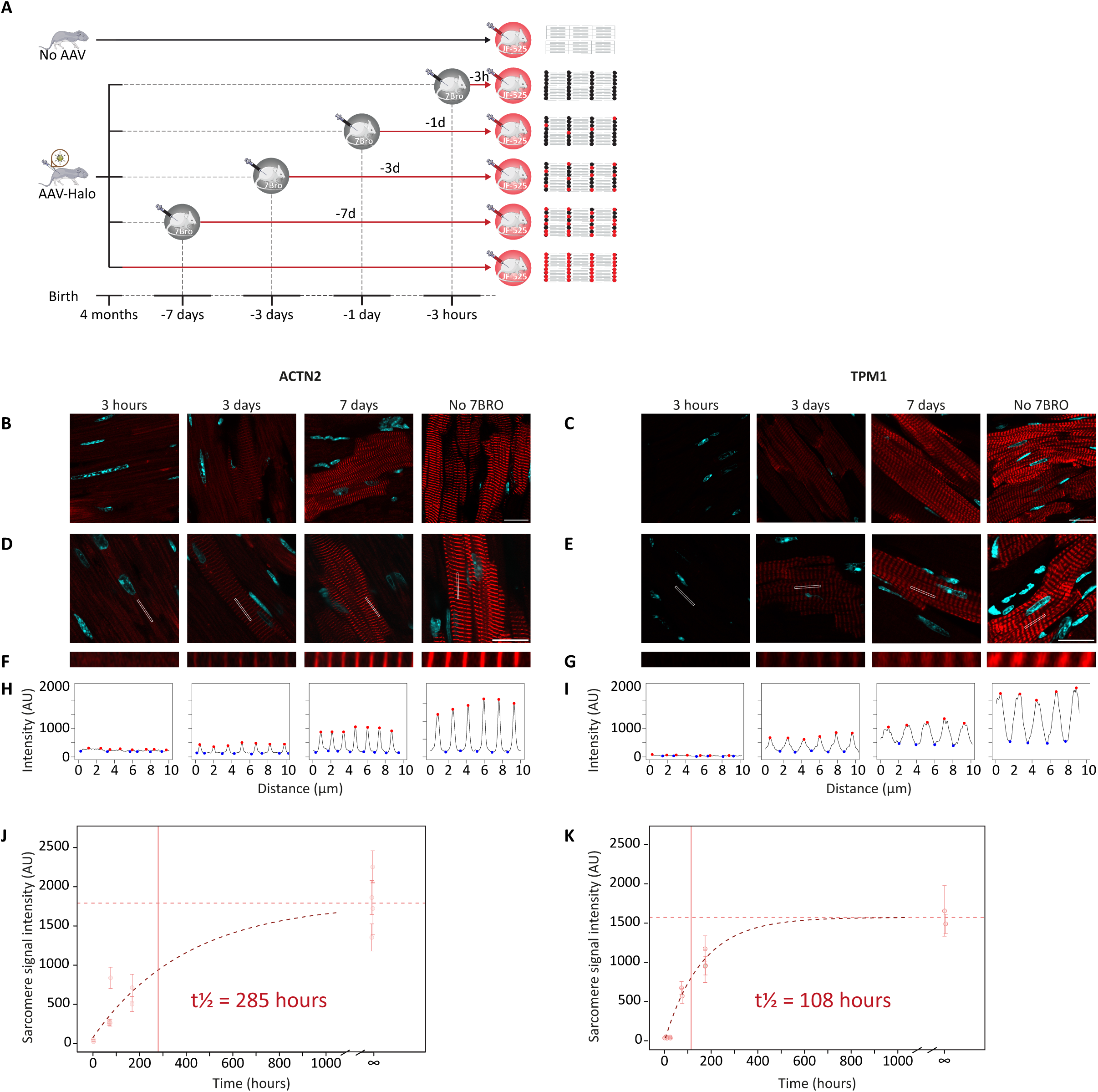
Imaging of sarcomeric protein turnover in vivo. **A,** Neonatal mice were divided into six groups: In the ‘No AAV’ group no AAV injection was administered, while in the other five groups mice received AAV9 encoding for either ACTN2-Halo or TPM1-Halo. In adulthood all groups received an intravenous injection of the JF-525 Halo-ligand, and their hearts were harvested for analysis after 40 minutes. Mice in the ‘No AAV’ and the ‘No 7BRO’ groups did not receive intravenous injection of the non-fluorescent blocker ligand 7BRO, while mice in the other groups received 7BRO 3 hours or 1, 3, or 7 days before the JF-525 ligand injection respectively. **B-G**, Representative images of the four groups of mice transduced with ACTN2-Halo (left) or TPM1-Halo (right) at low magnification (**B**,**C**, scale bar = 15µm), and higher magnification (**D**,**E**, scale bar = 10µm). A 10 µm region of interest along the myofibril is marked in (**D,E**) by a white rectangle and is shown at higher magnification in (**F,G**). **H-I**, Analysis of the JF-525 signal amplitude along the bar shown in **F,G**. The red and the blue dots are the local maxima and minima, identified automatically. **J-K**, The sarcomeric signal amplitude is plotted as a function of time after the 7BRO injection. Each dot represents the average of the sarcomere signal intensity from one mouse with ± SD whiskers. The ‘No AAV’ groups were regarded as time 0, the ‘No 7BRO’ as the maximal possible signal, marked by a horizontal red dashed line. A fitted 1 minus exponential curve is shown as a dark red dashed line and the calculated half-life as a red vertical line (n = 10 cells/mouse, from n = 2-4 mice/group for ACTN2-Halo; n=10 cells/mouse, from n = 2-3 mice/group for TPM1-Halo).

### Imaging endogenous TTN turnover in vivo

While AAVs are efficient at expressing Halo-tagged sarcomeric proteins, their limited cloning capacity and variability in transduction efficiency may restrict their use. We therefore again used the mice in which HaloTag was knocked-in the I-band section of the giant sarcomeric protein TTN^33^ to study the turnover of TTN in vivo. We employed a similar labeling approach to the one we used in the AAV transduced mice with injection of the non-fluorescent blocker 7BRO, followed by an injection of JF-525. We used similar negative controls with wild type mice that do not have Halo-tagged TTN and Halo-tagged TTN mice in which the fluorescent ligand was injected 3 hours after injection of the blocker, to confirm complete blocking of existing Halo-tagged TTN. We also injected Halo-ligand JF-525 to Halo-tagged TTN mice that did not receive the 7BRO blocker to measure the maximal possible labeling intensity. Here we included more labeling time points, with injection of Halo-ligand JF-525 at 3, 7, and 14 days after the injection of the blocker (Figure 5A). We also added Halo-ligand JF-525 to the cardiac sections to ensure complete labeling. Representative images of the sections and of cells at low and high magnification showed no sarcomeric signal with injection after 3 hours, indicating blockade was complete, and increasing sarcomeric signal at 3, 7, and 14 days (Figure 5B through 5D). The group that did not receive the non-fluorescent blocker showed maximal signal intensity. We used the same analysis approach we used for the AAV transduced hearts. We drew a region of interest along the myofibril axis on individual cardiomyocytes (Figure 5D) and used an automated tool to calculate the median amplitude of the sarcomeric signal along the line (Figure 5E). We fitted the data to a 1 minus exponential decay function and estimated the half-life of endogenous TTN to be 191 hours (Figure 5F). This number is more than 6-fold longer than the turnover of TTN in cultured NMVMs, indicating a marked slowdown in TTN turnover after maturation. To measure the half-life of TTN in neonatal mice in vivo, we injected 1 day old pups with the 7BRO blocker, and harvested the hearts after 3 hours, 1, 3, 7, or 10 days with labeling of Halo-TTN with JF-525. We also included Halo-tagged TTN mice that did not receive the 7BRO blocker to measure the maximal possible labeling intensity (Figure S10A). In neonatal mice, TTN has a half-life of 86 hours, implying a turnover rate more than double that of adult mice (Figure S10B through S10F).

**Figure 5.**
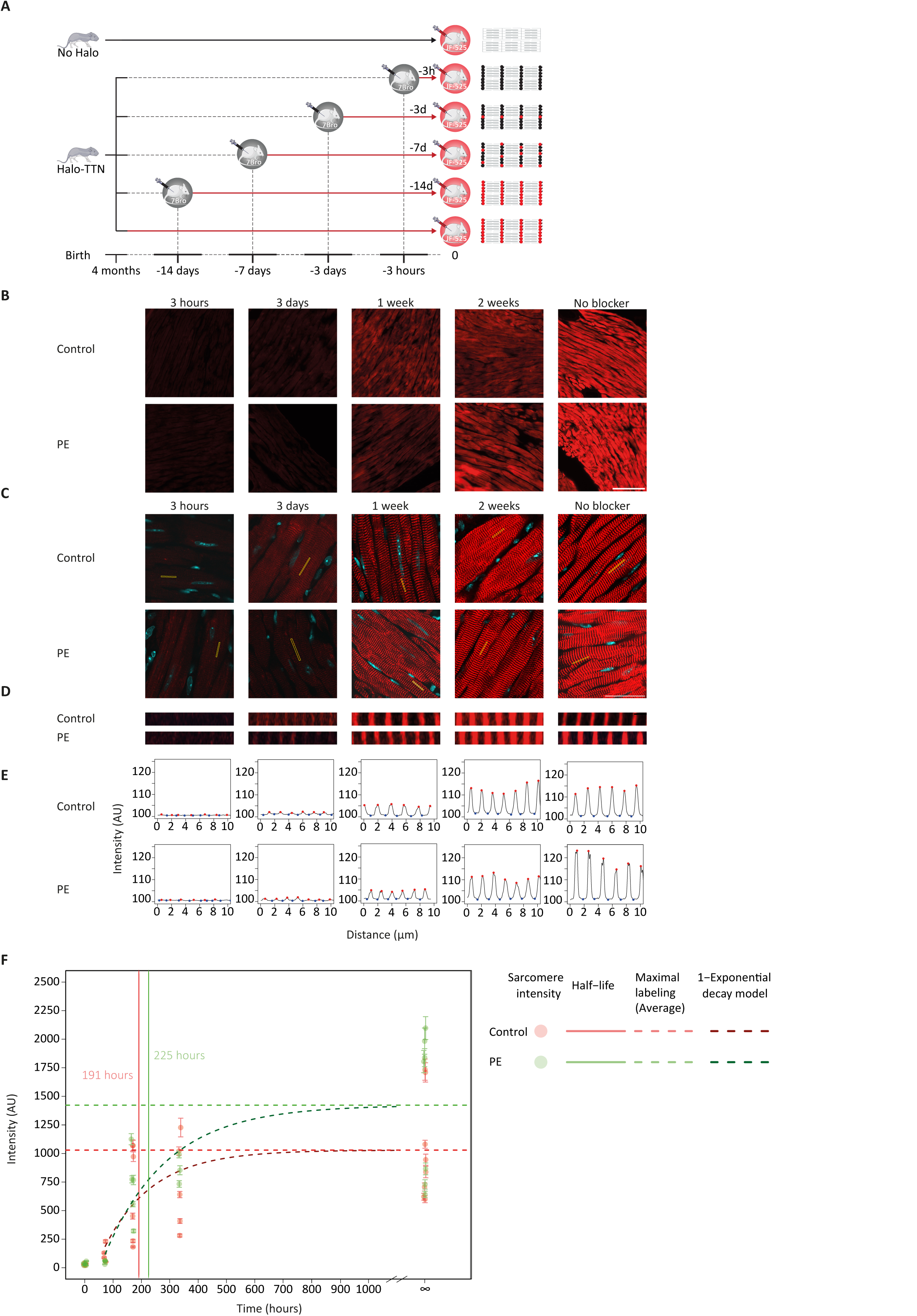
In-vivo turnover of endogenous TTN in the normal heart and in hypertrophy. **A,** Experiment design. Adult Halo-TTN knock-in mice received a non-fluorescent Halo-ligand (7BRO). After a chase period of 3 hours, 3 days, 1 week, or 2 weeks, mice received the JF-525 ligand, and the hearts were harvested. To measure the maximal labeling, one group of HaloTag-TTN knock-in mice received the fluorescent ligand without the 7BRO blocker. A negative control group of wild-type mice also received the fluorescent ligand without the 7BRO blocker. **B-D,** Representative images of cardiac sections from the different groups for control and phenylephrine (PE) treated mice at low (**B**, scale bar = 100µm), and high magnification (**C**, scale bar = 30µm). The yellow box in (**C**) indicates a region of interest of ∼10 µm length used for quantification and is shown at high magnification in (**D**). **E,** Quantification of signals from (**D**). The local maxima and minima were identified automatically and marked in red and blue dots, respectively. **F,** Quantification of sarcomere labeling intensity from control mice (red) and mice that received PE (green). Each dot represents the mean sarcomere intensity of 10 cells from 10 different fields, acquired from a section of a single mouse heart. The error bars represent the standard error. The no blocker control group data is indicated in infinity. The horizontal dashed line is the mean intensity in the no blocker positive control group and was used as total protein reference in the model. The curved dark dashed line is the fitting curve of the data to 1 minus exponential decay model. The vertical solid lines indicate the half-life of the TTN protein calculated from the model. Time is indicated in hours, and intensity in arbitrary units (AU). (n = 10 cells/mouse, from n = 3-8 mice/group, from N=4 independent experiments, ANOVA between control and PE p=0.205)

We next aimed to analyze the Halo-tagged TTN mice after induction of cardiac hypertrophy. Administration of phenylephrine (PE) activates α1-adrenergic receptors and is known to stimulate cardiomyocyte protein synthesis in mice.^40^ We used a PE injection protocol in the Halo-tagged TTN mice and verified that it indeed resulted in a significant increase in normalized heart weight, indicative of hypertrophy (Fig S11A). We asked whether hypertrophy and an increased protein synthesis would also result in increased turnover rate of TTN. We therefore included groups of mice in which we injected PE, in parallel with the Halo-ligand pulse chase protocol (Figure 5B through 5F and Fig S11B). The average maximal labeling intensity observed in the group that did not receive the 7BRO blocker was higher in the PE-treated mice, but the difference was not statistically significant. Accordingly, the half-life calculated for the PE group was slightly longer than in the control group (225 vs 191 hours), but the difference was not statistically significant. These data suggest that the turnover of titin is not significantly accelerated in hypertrophy. The use of global fitting for analysis of the data showed similar half-life values (Figure S12).

Finally, we analyzed the turnover rates in different regions of the heart. We chose the left ventricular free wall, the interventricular septum and the right ventricular free wall as representative regions and measured the half-life of TTN in each of them. Our analysis showed that TTN is turned over in different regions of the heart at similar rates (Figure S13).

## Discussion

Here we developed a pulse-chase approach based on the ability of Halo-tagged proteins to covalently bind fluorescent ligands. This approach allowed us to visualize pre-existing and newly translated proteins, measure their assembly and degradation rates, and uncover the mechanisms of sarcomeric maintenance. We showed that different sarcomeric proteins are uniformly turned over at similar rates, that turnover is slower in vivo in adult mice than in cultured neonatal cardiomyocytes or in neonatal mice, and that the turnover of the giant protein TTN is uniform throughout the heart. We challenged the prevailing ‘protein pool’ hypothesis for sarcomeric maintenance and found that assembly is unidirectional - newly translated proteins continuously and uniformly enter the structure, whereas already assembled proteins can only be removed and degraded. Further, we demonstrated that older and newer sarcomeric proteins are not distinguished in the degradation process, and that proteolytic extraction from the complex is a rate limiting step in the turnover.

Since expressed sarcomeric proteins stoichiometrically replace endogenous ones^41^ we could use virally expressed Halo-tagged sarcomeric proteins, and we confirmed that this applied to endogenous proteins as well. We relied on the TTN Halo-tag knock-in mice to image the turnover of the giant protein TTN. Since such tagged mice were not available for other proteins, we developed a simple HITI approach that can be used to knock-in HaloTag to endogenous proteins in cultured cardiomyocytes by expressing Cas9, the donor HaloTag cassette, and in-vitro transcribed sgRNAs. We used this simple approach to follow the replacement and turn-over of endogenous ACTN2 and MYOM1, and we believe that this approach can be adapted to almost any protein.

Proteomic approaches were used to measure the half-lives of cardiac proteins in cultured cells and in vivo.^42–44^ Coupled to mass-spectrometry, such analyses are powerful and allow the study of thousands of proteins in parallel. However, proteomics averages the half-life of proteins over the entire heart tissue and cannot differentiate between cardiomyocytes and non-cardiomyocytes. We did not measure the half-life of proteins per se, but rather calculated the turnover inside the sarcomeric structure in cardiomyocytes. Nevertheless, for the giant protein TTN the half-life of turnover measured by us, ∼8 days, was similar to the average 11.8 days half-life calculated by proteomics.^42,45,46^ Likewise, the half-lives we measured for ACTN2 and TPM1 of 11.9, and 4.5 days were comparable to the proteomics calculated half-lives of 11.4 and 7.15 days respectively ^42^. A hypertrophic stimulation did not alter the half-life of TTN, as was also observed with proteomics^42^. The main advantage of our approach, however, is that it allowed us to study and image the turnover dynamics of proteins in living cultured cardiomyocytes and in vivo at the single cell and even at the single sarcomere level. We used a first-order kinetics model to analyze the assembly and degradation of the protein independently, without assuming a steady state, and found that the half-life for assembly was longer than that calculated for the degradation. This difference could reflect a true decrease in the total content of the sarcomeric protein in the cardiomyocytes throughout the measurement time, but we believe that it is more likely a result of bleaching during time lapse imaging which accelerated the decrease in green fluorescence and delayed the accumulation of far-red fluorescence. Nevertheless, the relative comparisons between sarcomeres, cells, or conditions would not be affected, even if some corrections are needed to calculate the exact half-lives.

With our approach, we were able to understand the key principles underlying protein complex maintenance, specifically for sarcomeres. Although it has been widely reported that sarcomeric proteins could be recycled^13–15^, we showed that the maintenance is unidirectional - new proteins can get into the complex, while older ones can only exit, and cannot be recycled. Previous studies of sarcomere turnover used FRAP with genetically encoded fluorophores, typically eGFP or YFP, and extended the analysis time to many hours to show recovery. An inherent assumption of FRAP is that the transition between bright and dark states is irreversible, and that fluorescence recovery is only caused by fluorescent molecules diffusing into the bleached area. It is also possible, however, for fluorescent molecules to convert into a reversible dark state, in a process called photoswitching. Several fluorescent molecules, including GFP, were shown to exhibit photoswitching^47^, and such reversible photoswitching may appear as mimicking diffusion. Indeed, one study that used fluorescently tagged TTN found extensive reversible photobleaching that accounted for most of the TTN fluorescence recovery.^48^ Our experiments did not reveal any recovery of OG Halo-tagged sarcomeric proteins, even when we bleached at a submaximal level. This fluor is likely less prone to photoswitching and unlike genetically encoded fluorophores does not need to fold or mature. We performed experiments in fibroblasts, where expressed sarcomeric proteins can freely diffuse to show that recovery following FRAP can be detected in our imaging setting. We did observe some quick recovery for MYL2, suggesting there may be a small diffusible pool for this protein.

In contrast with assembly that is ordered, degradation appeared to be stochastic. Cardiomyocytes degrade old and new proteins at similar rates, and likely cannot distinguish between them. We further showed that pharmacologic blockade of both calpains and proteasomal proteases delays the removal of existing proteins from the sarcomeric structure, indicating that proteolytic cleavage of sarcomeric proteins is required for their extraction. The removal of existing proteins, however, was slowed, but not completely stopped. We assume that the incomplete response to these inhibitors is due to incomplete protease blockade, as we were limited in the dose due to toxicity, but we cannot exclude that a certain portion of sarcomeric proteins can exit the structure without being cleaved first. Our result suggests that calpain mediated cleavage is the rate limiting step, but due to the potential limited specificity of pharmacological inhibitors further genetic studies are needed to specifically delineate the roles of these proteolytic systems. These data, together with the observation that the turnover of over-expressed ACTN2 occurs at the same rate as that of the endogenous protein, implies that the proteolytic extraction from the sarcomere is a rate limiting step in the turnover of sarcomeric proteins. These findings also suggest that the degradation of sarcomeric proteins may be regulated. Furthermore, the proteolytic requirement for sarcomeric protein replacement we show supports the proteasome functional insufficiency hypothesis of heart failure.^49,50^

There are multiple sarcomeric proteins, many with different isoforms. The turnover principles we identified were similar for all the sarcomeric proteins we studied. Due to the complexity of our studies, such as the need for long and stable imaging of 72 hours for each cell, we were unable to extend the study to additional sarcomeric proteins. As these proteins all belong to the same complex it is highly likely that they share the same turnover mechanisms; however, we cannot rule out the possibility that some exceptions exist.

In conclusion, we discovered mechanisms of sarcomere maintenance that could be useful for developing future treatments for genetic cardiomyopathies as well as for heart failure and hypertrophy. In particular, the requirements for enzymatic cleavage as a rate limiting step in the process of replacement may be exploited to increase or delay the turnover of proteins in the sarcomeric structure. Finally, while we demonstrated how sarcomeres are maintained in cardiomyocytes, our findings likely apply also to other heteromeric complexes in other cells.

## Non-standard Abbreviations and Acronyms

FRAP: Fluorescence recovery after photobleaching
NRVMs: Neonatal rat ventricular cardiomyocytes
OG: Oregon Green
7BRO: 7-bromoheptanol
JF-635: Janelia Fluor 635
NMVMs: Neonatal mice ventricular cardiomyocytes
HITI: homology-independent targeted integration
sgRNA: single guide RNA
AAVs: adeno associated viruse
JF-525: Janelia Fluor 525
ALLN: N-acetyl-leucyl-leucyl-norleucinal
PE: phenylephrine

## Acknowledgements

The authors wish to thank the Biomedical Core Facility at the Faculty of Medicine, Technion, the Technion Genomic Center, and the Pre – Clinical Research Authority at the Technion. The authors wish to thank **Luke D Lavis,** Janelia Research Campus, Howard Hughes Medical Institute, Ashburn, Virginia, USA for his generosity in providing the Janelia Fluor HaloTag ligands.

## Supplementary materials

Supplementary methods

Supplementary figures S1-S13

## Figure legends

**Figure S1. Models of sarcomere maintenance.** Schematic illustration of the photobleach pulse-chase experiment and the possible outcomes according to two sarcomere maintenance models. Halo tagged proteins are expressed using adenoviral vector and labeled with green fluorescent ligand, that is bleached in a small square area. The cardiomyocytes are then incubated with a far-red ligand. **A**, The protein pool model predicts the existence of a cytoplasmic pool of sarcomeric proteins composed of both recycled (green labeled) proteins and newly translated (far red labeled) proteins. Since the photobleached proteins are expected to recycle proteins with this pool, the bleached area is expected to fill with both green and red labeled proteins. **B**, The ordered assembly model on the other hand predicts that only new proteins can enter the structure and therefore the bleached proteins will only be replace by far-red labeled ones.

**Figure S2. Recovery of Oregon Green labeled ACTN2-Halo fluorescence in fibroblasts after bleaching A,** Representative images from a FRAP experiment performed in fibroblasts transduced with ACTN2-Halo and labelled with Oregon green. Images show the same fibroblast before the bleach (pre-bleach), and at 0,1, and 18 minutes after the bleach, with almost complete recovery of the green signal within 1 minute. The dashed white square is the photobleached field. The dotted yellow rectangle is the region of interest used for quantification. Scale bar = 10 µm. **B**, Quantification of the green florescence recovery after photobleaching. We calculated the photobleaching rate (r) by comparing the fluorescence of the non-frapped region of the cell before (Fc0) and after photobleaching (Fc). 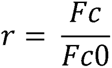 Then, we normalized the fluorescence intensity of the photobleached area by dividing its raw intensity (Fs) by the photobleaching rate (r). 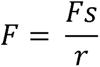

**Figure S3. Semi-automated image analysis of sarcomere turnover pipeline A,** representative image of pulse-chase experiment of cardiomyocytes transduced with Halo-tagged MYOM1, pre and after photobleach, over 72 hours. The image is a merge of Oregon Green signal of pre-exiting proteins, and JF-635 ligand far-red signal of newly translated proteins from Figure 1. The numbers above indicate the time in hours after the photobleach. The dashed white square indicates the bleached area. The yellow rectangles labeled **B** & **C** are the region of interest (ROI) chosen for the quantification. Scale bar = 20 µm. **B-C,** Fluorescent intensity of the ROI outside and inside the photobleached area (**B & C** respectively) is shown in green for the pre-exiting proteins, and in red for the newly translated proteins. Each graph represents one timepoint according to the representative image above. The distance is shown in µm, and the signal intensity is shown in arbitrary units (AU). The peaks and the nadir points were identified using a moving window of 1 µm, and labeled by blue and turquoise dots respectively. **D,** Example of analysis of a single timepoint signal. The grey rectangles represent the difference between a nadir and the adjacent peak. The yellow rectangles represent the difference between a peak and the adjacent nadir point. The median of these differences was used as the representative value for the specific timepoint and channel.

**Figure S4. Turnover rates in single sarcomeres are similar.** Single sarcomeric units from one myofibril in the unbleached region were individually analyzed using time lapse imaging to measure the decay of their green fluorescence intensity and to calculate their individual half-lives. The measured sarcomeric fluorescent intensity is shown on the left as dots, with fitted exponential decay (dotted lines) and calculated half-lives (bold dot) for each sarcomeric unit. Numbers show the range of calculated half-lives for the different sarcomeric units. Time lapse images are shown on the right, scale bar = 2µm. Two individual myofibrils, each containing several sarcomeric units are shown for each Halo-tagged sarcomeric protein: **A-B** ACTN2-Halo, **C-D** MYOM1-Halo, **E-F** Halo-MYL2, **G-H** Halo-MYBPC3. The analysis was extended beyond 72 hours in myofibrils where there was no significant movement. The images were taken every 12 hours instead of every 24 hours in some cells where the initial signal intensity was very strong.

**Figure S5. Validation of ACTN2 half-life measurements. A-C**, As an alternative to the live-imaging approach, NRVMs were transduced with ACTN2-Halo adenoviral vector. After 48 hours, OG ligand was added to the media, followed by 7BRO blocker ligand and medium replacement. After 0, 24, or 48 hours cells were incubated with TMRDirect Halo red ligand, and harvested for protein extraction. The cell lysate was analyzed by SDS-PAGE with in-gel measurements of fluorescence (**A**). The gel fluorescence in the green (top), red (middle), and merged (bottom) at the size of 140 kDa, corresponding to ACTN2-Halo shows decreasing green and increasing red fluorescence (**B**). Normalized fluorescence signal at 140 kDa is shown for the green and red channels with fitted exponential decay and accumulation curves (dotted lines) and calculated half-lives (bold dot and number), which are similar to the values measured by imaging (**C**). N = 2 independent replicates, with n= 3-4 culture plates per time point. **D-E**, As another alternative approach to assess the half-life of ACTN2, NRVMs were incubated with the protein translation inhibitor Cycloheximide (CHX) or control for 8 hours. Western blot analysis for ACTN2 showed decreased levels after blockade of translation (**D**) Quantification of the data shows significantly reduced ACTN2 levels and allowed estimation of the half-life of 26 hours for ACTN2 (**E**). N = 4 independent replicates with n = 6-7 culture plates.

**Figure S6. HITI to knock-in HaloTag into the C-terminus endogenous genes. A**, Illustration of the HITI targeting approach: a Cas9 site (sgRNA target) that is followed by a PAM (NGG) site was identified in the C-terminal end of *Actn2* before the stop codon. The scissors show the predicted cut site (top). A donor cassette encoding the C-terminal end of *Actn2* with no stop codon, followed by in-frame HaloTag, and flanked by the same Cas9 sites (sgRNA target) motifs was cloned into an AAV vector (middle). The direction of the sgRNA target sites flanking the donor cassette are inversed relative to the direction of the site in the genome. Integration of the cassette in the correct orientation to the genomic locus results in a chimeric junction that is no longer recognized by the sgRNA (bottom). Insertion in the inverse orientation on the other hand would re-create the sgRNA site and will be cut out again (not shown). **B**, Representative images from Cas9 expressing mouse cardiomyocytes transduced with Actn2-HaloTag donor cassette virus and with either control non-targeting sgRNA (left) or *Actn2* c-terminus sgRNA (right) encoding virus after addition of the Oregon Green Halo-Ligand (time = 0 hours) showing cross-striated pattern only in cardiomyocytes transduced with both donor and *Actn2* sgRNA. A 10 µm linear region of interest drawn along the myofibril is marked by a yellow rectangle. **C**, The green -fluorescence intensity along the linear region of interest drawn in **B**. **D-E,** Cas9 expressing NMVMs were targeted by HITI to knock-in Halo to the C-terminus of *Actn2* (HITI) or over-expressed with ACTN2-Halo virus (OE). SDS-PAGE analysis of fluorescence showed much higher levels of ACTN2-Halo with over-expression by transduction, compared with knock-in. **F**, Representative time-lapse images of NMVMs after either HITI knock-in of HaloTag to the C-terminus of endogenous *Myom1* gene (endogenous), or after over-expression with Adenoviral vector encoding for N-terminal fragment of MYOM1-Halo (exogenous). Cells were pulsed with JF-635 ligand and imaged at 0, 24, 48, and 72 hours. Nuclear aggregate were observed only with the over-expressed N-terminal fragment of MYOM1-Halo (exogenous). Scale bar = 15µm. **G**, Normalized sarcomeric fluorescence amplitude with fitted exponential decay curve (dotted lines) and calculated half-life (bold dot and number; N=21-15 cells for endogenous and exogenous groups from N=3 independent experiments) showing similar half-life for the full length endogenous MYOM1-Halo (pink) and the over-expressed exogenous N-terminal fragment of MYOM1-Halo.

**Figure S7. Alternative protocol to assess degradation rate of new and old proteins. A**, Very high R^2^ values for fitting of the data in Figure 3C showing excellent fit with exponential curves. **B**, Alternative degradation experiment timeline - Cardiomyocytes expressing MYOM1-Halo for 48 hours were incubated with the OG ligand followed by blocking with 7BRO for 12 hours, washing, and then with the JF-635 Halo-ligands for an additional 24 hours ensuring the youngest OG labeled proteins are at least 12 hours older than the oldest JF-635 labeled proteins, and imaged in time-lapse for 72 hours. **C,** Representative images of cardiomyocytes at 0,24, 48, and 72 hours show a gradual decline of the sarcomeric signal in both the green and far-red channels. Scale bar = 10.25µm. **D-E,** Quantification of normalized sarcomeric fluorescence intensity in the green (OG) and far-red (JF) channels (**D**), showing the half-lives of newer and older proteins in individual cardiomyocytes did not differ (**E**) (n=32 cells, N=3 independent experiments, two tailed paired Student’s t-test p=0.63).

**Figure S8. Proteolytic cleavage determines the turn-over rate of sarcomeric proteins. A**, Western blot showing an increase in ubiquitinated proteins following MG-132 treatment for 24 hours. **B-C**, Western blot and quantification showing a significant increase in ACTN2 level following MG-132 treatment for 24 hours, which corresponds to a half-life of 21.4 hours. N = 2 independent replicates with n = 3-4 culture plates. **D-F**, Representative images (**D**) and quantification (**E-F**) of Halo-TTN in knock-in cardiomyocytes labeled with OG ligand and followed for 72 hours. Normalized sarcomeric fluorescence intensity for control (light green, Ctrl) and ALLN treated (dark green, ALLN) (**E-F**) shows a significantly longer half-life of Halo-TTN in ALLN-treated than in control cardiomyocytes (n=33-38 cells, N=3 independent experiments, two tailed Student’s t-test p<0.0005).

**Figure S9. Inhibition of the proteasome results in aggregation of over-expressed and of endogenous sarcomeric proteins. A-B**, Representative images of cardiomyocytes overexpressing ACTN2-Halo (**A**) or with Halo-TTN knock-in (**B**) treated with control or MG-132 for 0,24,48, or 72 hours as in Figure 3K and 3L in the green (pre-existing), far-red (newly-synthesized) or inset with merged green and far-red channel (dashed square). For both the over-expressed ACTN2-Halo and the endogenous Halo-TTN we see integration of newly synthesized far-red labeled proteins to the sarcomere in a cross striated patten as well as multiple aggregates of newly translated proteins (yellow dotted circle). Scale bar = 10µm.

**Figure S10. In-vivo turnover of endogenous TTN in the neonatal heart in vivo A,** Experiment design. Neonatal Halo-TTN knock-in mice received a non-fluorescent Halo-ligand (7BRO). After a chase period of 3 hours, 1,3,7, or 10 days mice received the JF-525 ligand. To measure the maximal labeling, one group of HaloTag-TTN knock-in mice received the fluorescent ligand without the 7BRO blocker (No 7BRO). **B-D,** Representative images of cardiac sections from the different groups at low (**B**, scale bar = 15µm), and high magnification (**C**, scale bar = 10µm). The white box in (**C**) indicates a region of interest of ∼10 µm length used for quantification and is shown at high magnification in (**D**). **E,** Quantification of signals from (**D**). The local maxima and minima were identified automatically and marked in red and blue dots, respectively. **F,** Quantification of sarcomere labeling intensity. Each dot represents the mean sarcomere intensity of 10 cells from 10 different fields, acquired from a section of a single mouse heart. The error bars represent the standard error. The no blocker control group data is indicated in infinity. The horizontal dashed line is the mean intensity in the no blocker positive control group and was used as total protein reference in the model. The curved dark dashed line is the fitting curve of the data to 1 minus exponential decay model. The vertical solid lines indicate the half-life of the TTN protein calculated from the model. Time is indicated in hours, and intensity in arbitrary units (AU). The half-life of TTN measured for adult mice was more than doubled the half-life measured in neonatal mice (n = 24-62 cells/mouse, from n = 3-8 mice/group, from N=12 independent experiments)

**Figure S11. Phenylephrine induced hypertrophy model. A**, A composite bar chart (mean ± standard error) with induvial values of heart weight normalized to body weight ratio (HW/BW in mg/g) of titin (TTN)-HaloTag knock-in mice, which did or did not receive phenylephrine (PE) 10 mg/kg in three doses over 7 days. (n=8-10 mice per group, * Student’s t-test p-value =0.000043). **B**, The PE injection protocol we used for the TTN-HaloTag knock-in mice, where mice received 3 injections in the last 7 days of experiment.

**Figure S12. Global fitting to analyze TTN half-life.** In addition to the fitting approach used in Figure 6 to evaluate Halo-TTN half-life in control and hypertrophic hearts we used global fitting. **A**, Data analysis of TTN half-life using global fitting. **B**, Comparison of the calculated half-life of TTN in control and hypertrophic hearts derived from local and global fitting, showing very small change.

**Figure S13. Titin is turned over at similar rates in different regions of the heart. A**, We separately calculated the turnover of titin (TTN) in three cardiac regions: the left ventricular free wall, the interventricular septum and right ventricular free wall. Representative cardiac sections from these regions are shown, left scale bar = 805 µm, right scale bar = 32 µm. **B**, The sarcomeric signal intensity is plotted as a function of time from the 7BRO blocker injection, and each dot represents the mean of the sarcomere signal intensity from one mouse. A fitted 1-exponential decay curve is shown as a dashed line, and the calculated half-life as a solid vertical line. The dots, horizontal dashed lines, solid lines, and turnover-half lives are colored by cardiac region. Half-life of TTN was similar in all regions. (n = 7-25 cells/mouse/region, from n = 1-7 mice/timepoint).

