## Supplemental Figures S1-S13 for "Imaging of existing and newly translated proteins elucidates mechanisms of sarcomere turnover"

Figure S1

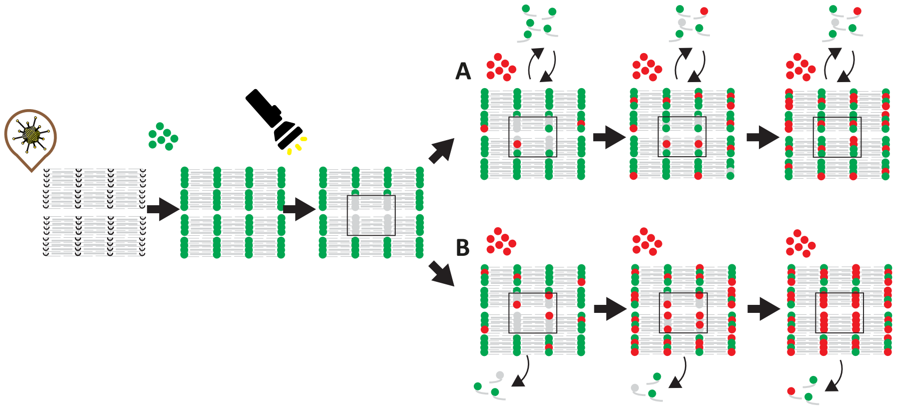

Figure S2

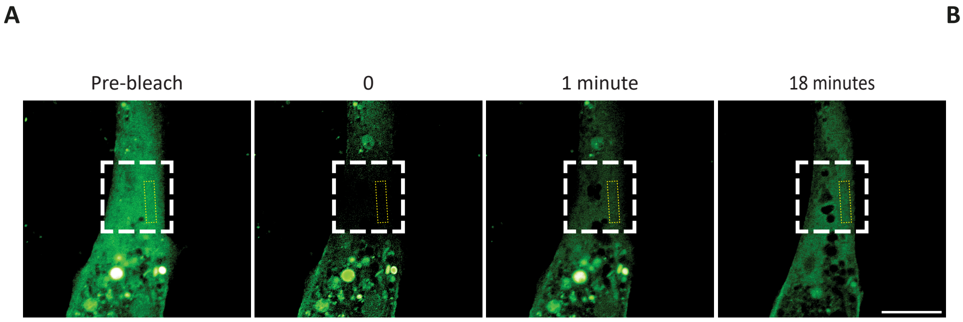

A
Pre-bleach
0
1 minute
18 minutes


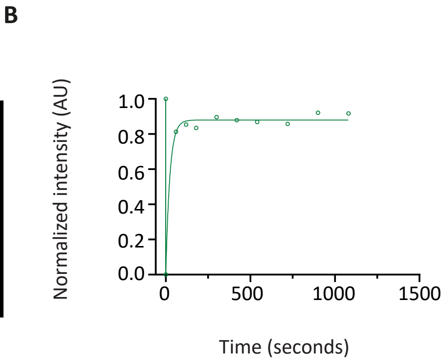

B
Normalized intensity (AU)
1.0
0.8
0.6
0.4
0.2
0.0
0
500
1000
1500
Time (seconds)

Figure S3

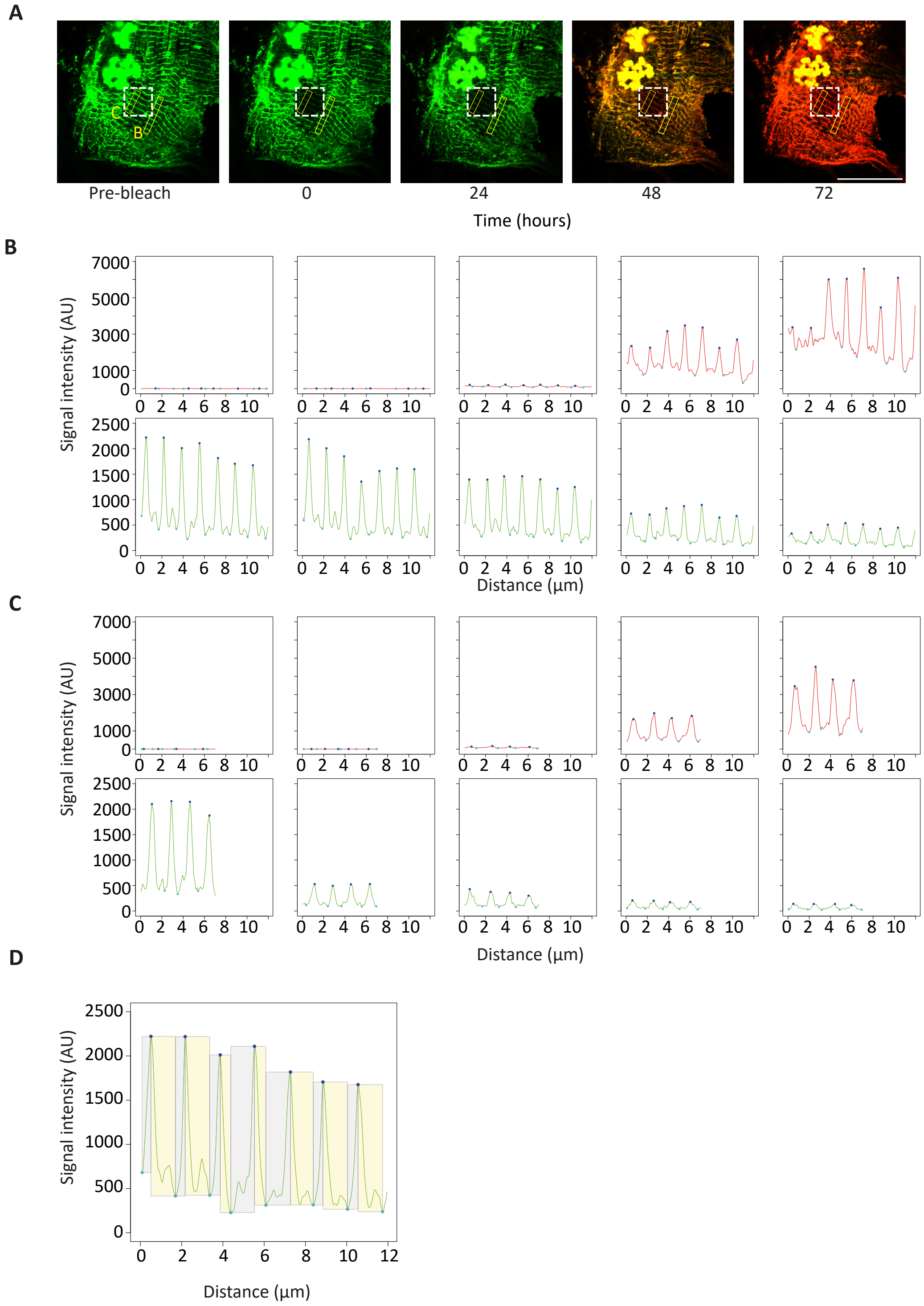
A
C
B
Pre-bleach
0
24
48
72
Time (hours)
B
Signal intensity (AU)
Distance (µm)
C
Signal intensity (AU)
Distance (µm)
D
Signal intensity (AU)
Distance (µm)

### Figure S4

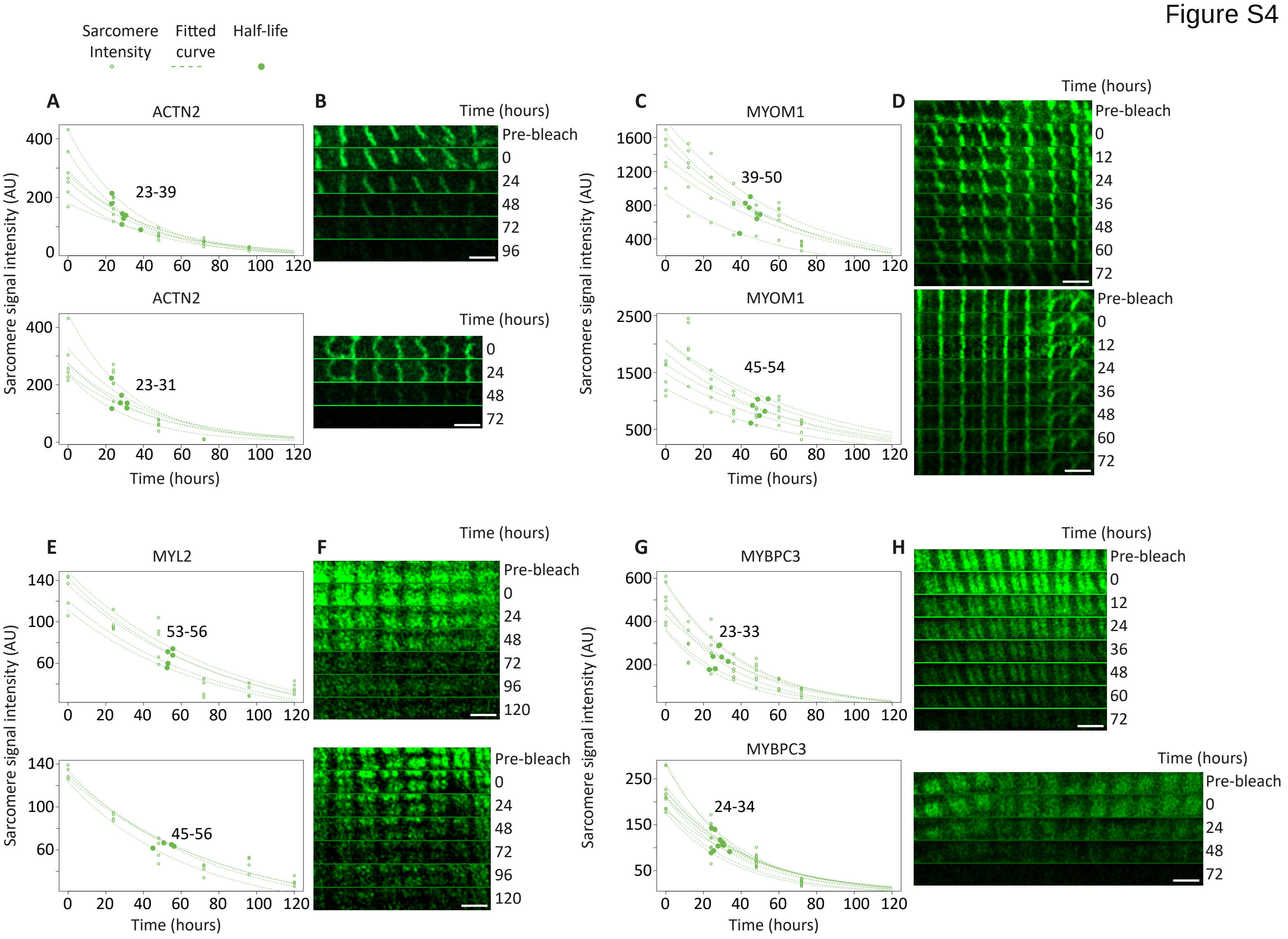

Sarcomere Intensity
Fitted curve
Half-life
A
ACTN2
23-39
ACTN2
23-31
Sarcomere signal intensity (AU)
Time (hours)
B
Time (hours)
Pre-bleach
0
24
48
72
96
Time (hours)
0
24
48
72
C
MYOM1
39-50
MYOM1
45-54
Sarcomere signal intensity (AU)
Time (hours)
D
Time (hours)
Pre-bleach
0
12
24
36
48
60
72
Pre-bleach
0
12
24
36
48
60
72
E
MYL2
53-56
45-56
Sarcomere signal intensity (AU)
Time (hours)
F
Time (hours)
Pre-bleach
0
24
48
72
96
120
Pre-bleach
0
24
48
72
96
120
G
MYBPC3
23-33
MYBPC3
24-34
Sarcomere signal intensity (AU)
Time (hours)
H
Time (hours)
Pre-bleach
0
12
24
36
48
60
72
Time (hours)
Pre-bleach
0
24
48
72

Figure S5

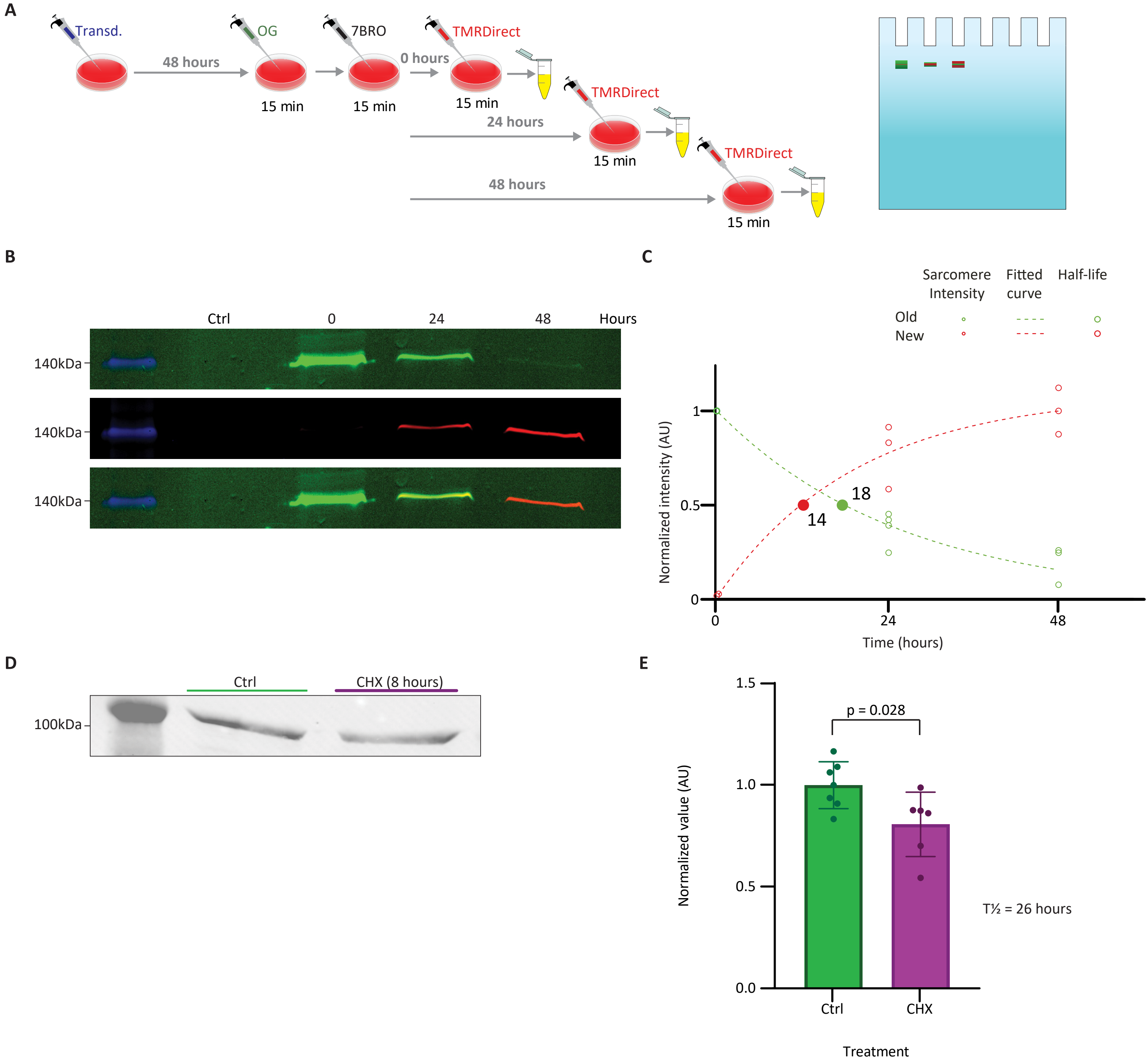
A
Transd.
48 hours
OG
15 min
7BRO
15 min
0 hours
TMRDirect
15 min
24 hours
TMRDirect
15 min
48 hours
TMRDirect
15 min
B
Ctrl
0
24
48
Hours
140kDa
140kDa
140kDa
C
Sarcomere Intensity
Fitted curve
Half-life
Old
New
Normalized intensity (AU)
1
0.5
0
18
14
0
24
48
Time (hours)
D
Ctrl
CHX (8 hours)
100kDa
E
p = 0.028
Normalized value (AU)
1.5
1.0
0.5
0.0
Ctrl
CHX
Treatment
T½ = 26 hours

Figure S6

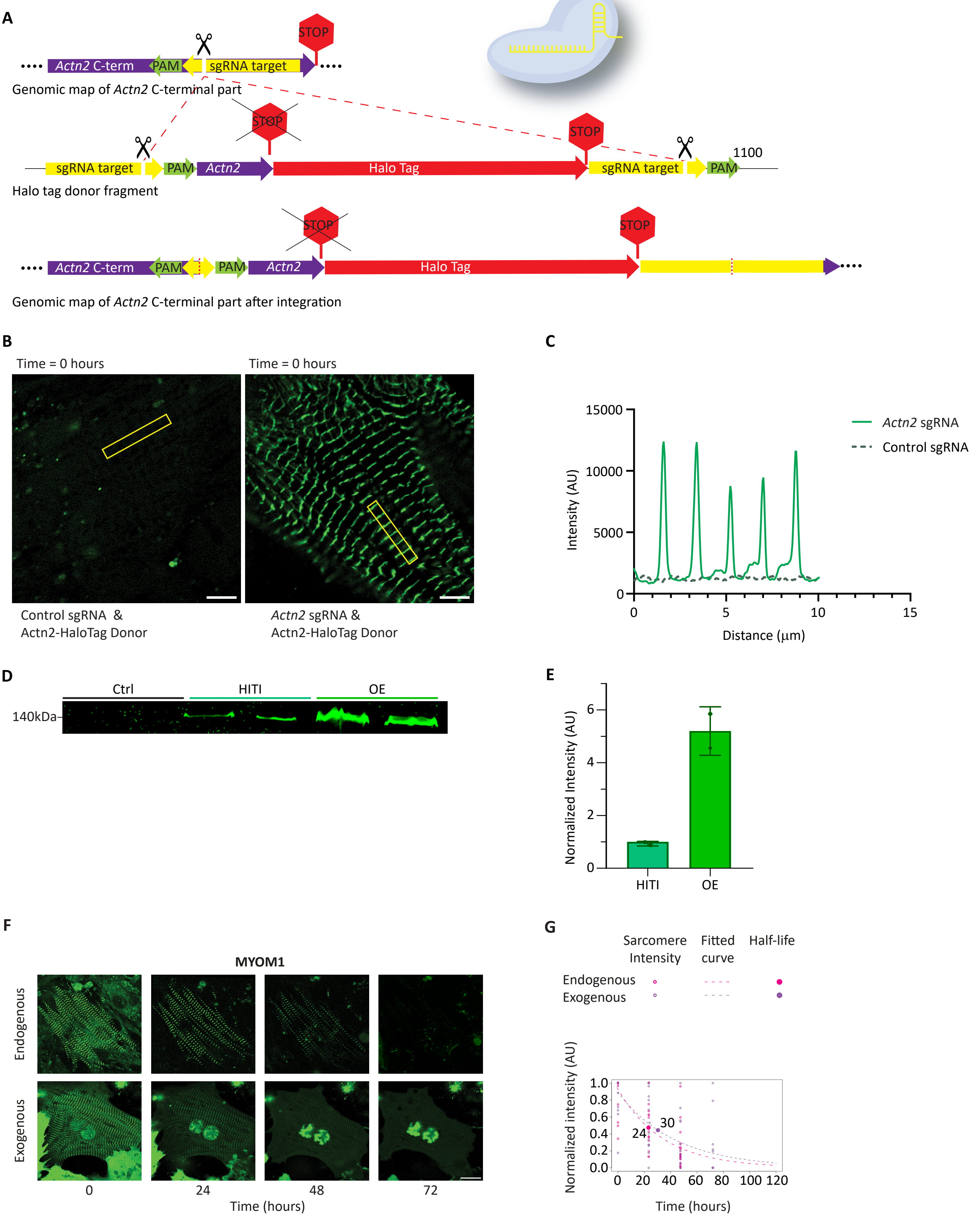

A
STOP
Actn2 C-term
PAM
sgRNA target
Genomic map of Actn2 C-terminal part
STOP
STOP
sgRNA target
PAM
Actn2
Halo Tag
sgRNA target
PAM
1100
Halo tag donor fragment
STOP
STOP
Actn2 C-term
PAM
PAM
Actn2
Halo Tag
Genomic map of Actn2 C-terminal part after integration
B
Time = 0 hours
Time = 0 hours
Control sgRNA &
Actn2-HaloTag Donor
Actn2 sgRNA &
Actn2-HaloTag Donor
C
Actn2 sgRNA
Control sgRNA
Intensity (AU)
15000
10000
5000
0
0
5
10
15
Distance (µm)
D
Ctrl
HITI
OE
140kDa
E
Normalized Intensity (AU)
6
4
2
0
HITI
OE
F
MYOM1
Endogenous
Exogenous
0
24
48
72
Time (hours)
G
Sarcomere Intensity
Fitted curve
Half-life
Endogenous
Exogenous
Normalized intensity (AU)
1.0
0.8
0.6
0.4
0.2
0.0
24
30
0
20
40
60
80
100
120
Time (hours)

Figure S7

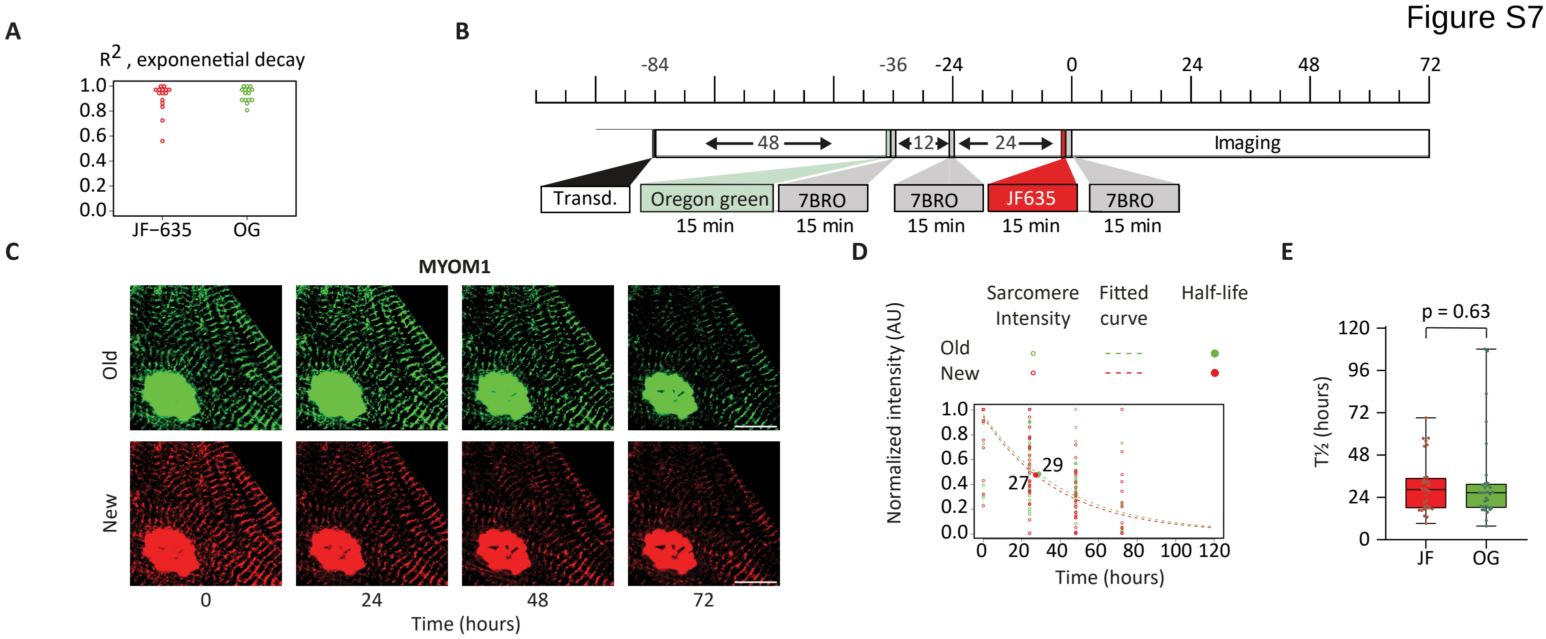

A
$R^2$, exponenetial decay
1.0
0.8
0.6
0.4
0.2
0.0
JF−635
OG
B
-84
-36
-24
0
24
48
72
48
12
24
Imaging
Transd.
Oregon green
15 min
7BRO
15 min
7BRO
15 min
JF635
15 min
7BRO
15 min
C
MYOM1
Old
New
0
24
48
72
Time (hours)
D
Sarcomere Intensity
Fitted curve
Half-life
Old
New
Normalized intensity (AU)
29
27
0
20
40
60
80
100
120
Time (hours)
E
p = 0.63
120
96
72
48
24
0
T½ (hours)
JF
OG

Figure S8

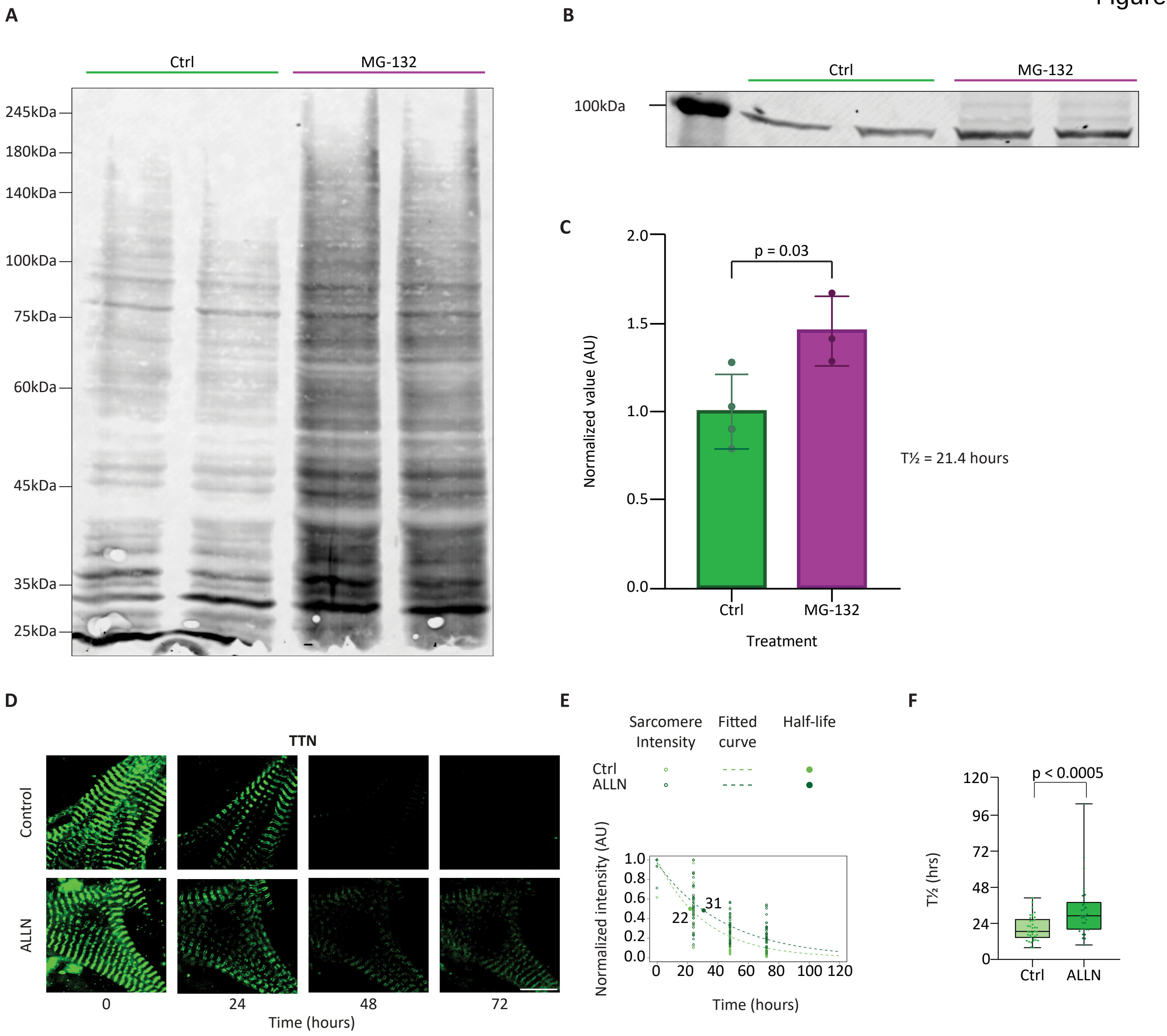
A
Ctrl
MG-132
245kDa
180kDa
140kDa
100kDa
75kDa
60kDa
45kDa
35kDa
25kDa
B
Ctrl
MG-132
100kDa
C
p = 0.03
Normalized value (AU)
2.0
1.5
1.0
0.5
0.0
Ctrl
MG-132
Treatment
T½ = 21.4 hours
D
TTN
Control
ALLN
0
24
48
72
Time (hours)
E
Sarcomere Intensity
Fitted curve
Half-life
Ctrl
ALLN
Normalized intensity (AU)
22
31
Time (hours)
F
p < 0.0005
T½ (hrs)
120
96
72
48
24
0
Ctrl
ALLN

Figure S9

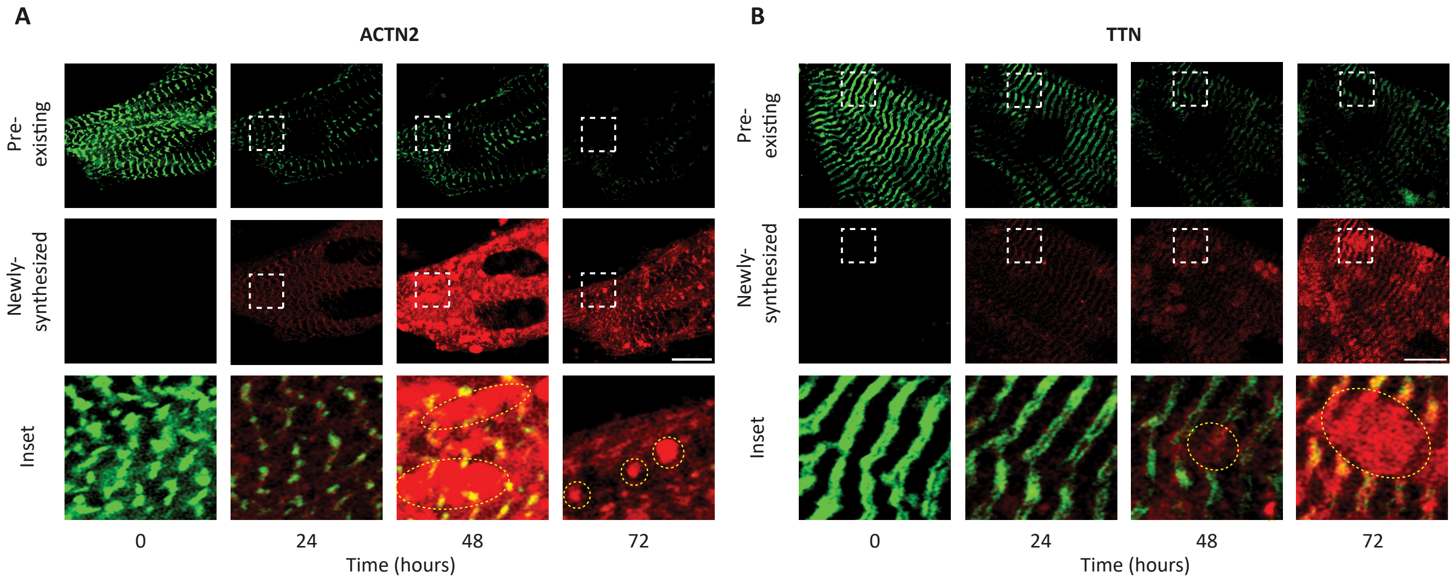
A
ACTN2
Pre-existing
Newly-synthesized
Inset
0
24
48
72
Time (hours)
B
TTN
Pre-existing
Newly-synthesized
Inset
0
24
48
72
Time (hours)

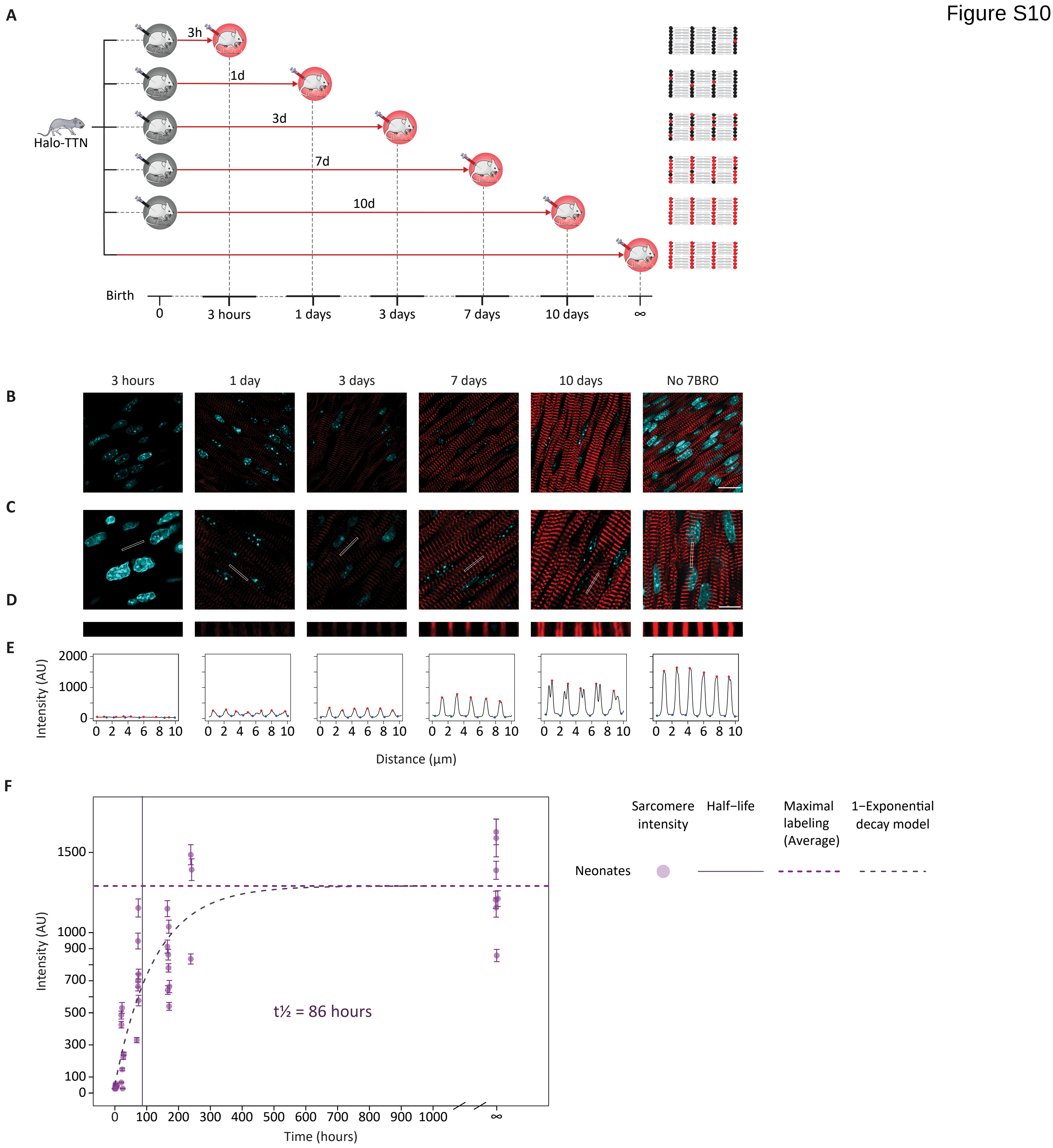

Figure S10
A
Halo-TTN
3h
1d
3d
7d
10d
Birth
0
3 hours
1 days
3 days
7 days
10 days
∞
B
C
D
E
3 hours
1 day
3 days
7 days
10 days
No 7BRO
Intensity (AU)
2000
1000
0
0 2 4 6 8 10
Distance (µm)
F
Intensity (AU)
1500
1000
900
700
500
300
100
0
t½ = 86 hours
0 100 200 300 400 500 600 700 800 900 1000 ∞
Time (hours)
Sarcomere intensity
Half−life
Maximal labeling (Average)
1−Exponential decay model
Neonates

Figure S11

A

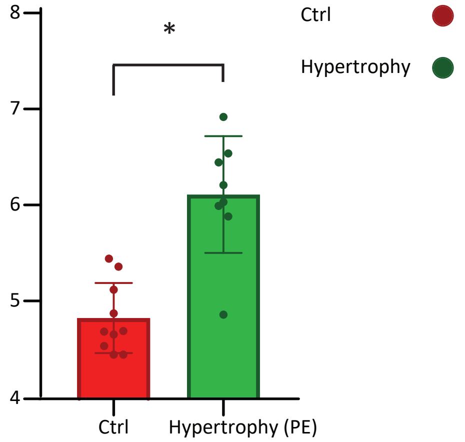


B

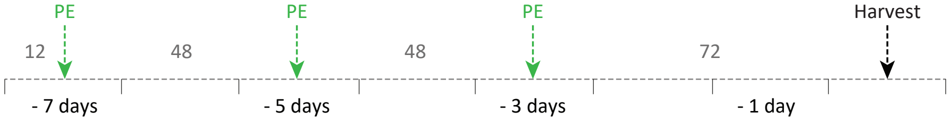

Figure S12

A

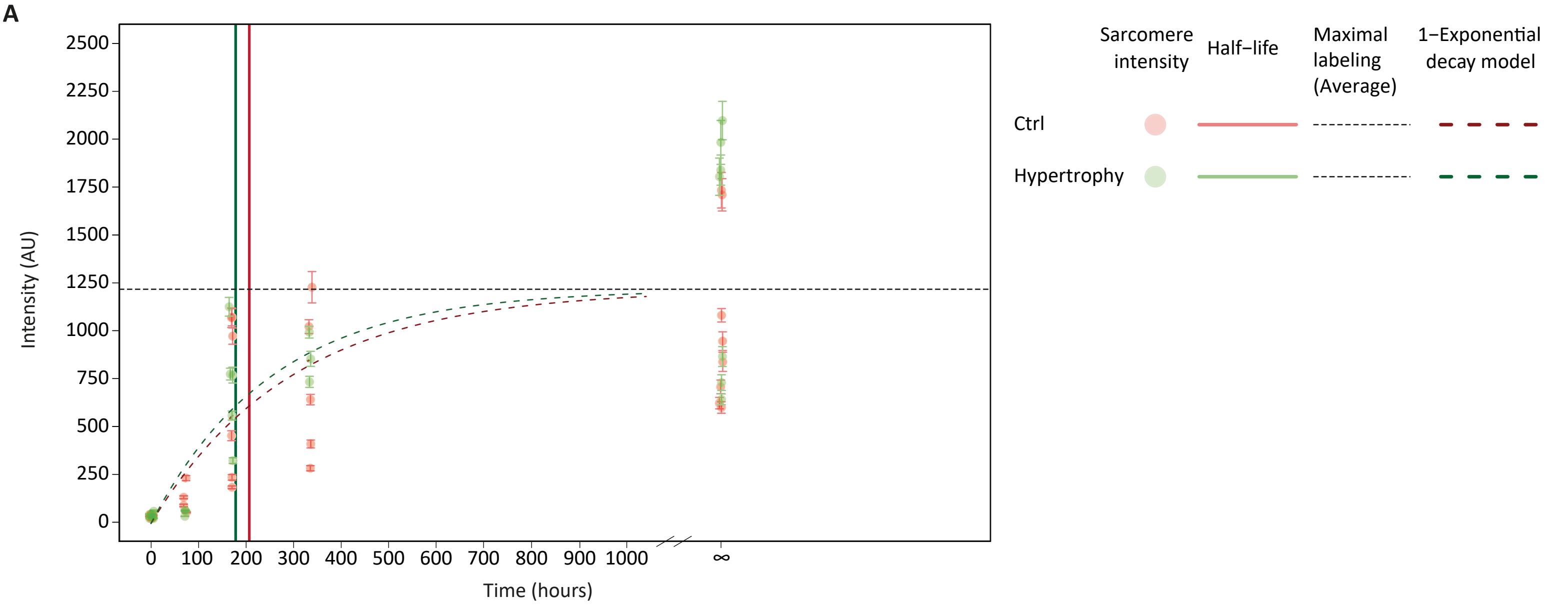


B

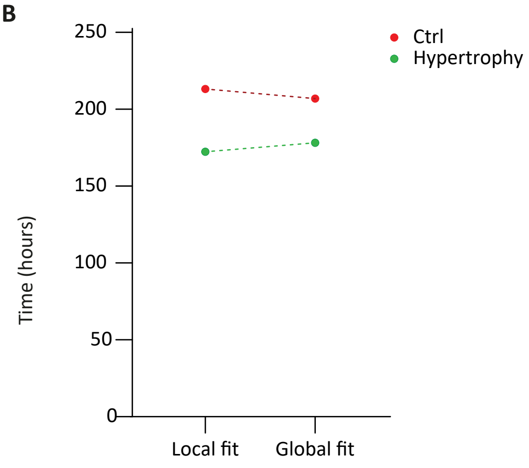

A

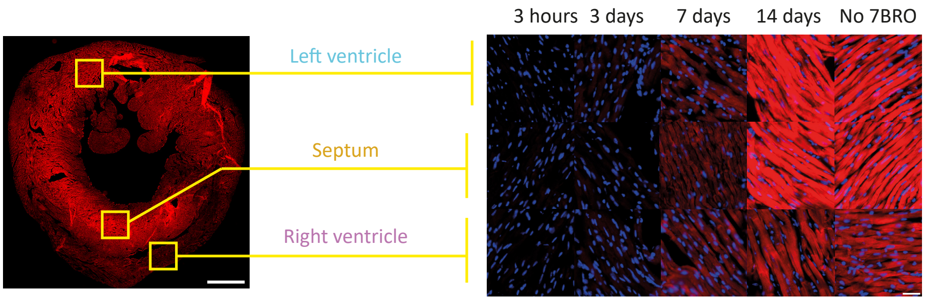
3 hours
3 days
7 days
14 days
No 7BRO
Left ventricle
Septum
Right ventricle

B

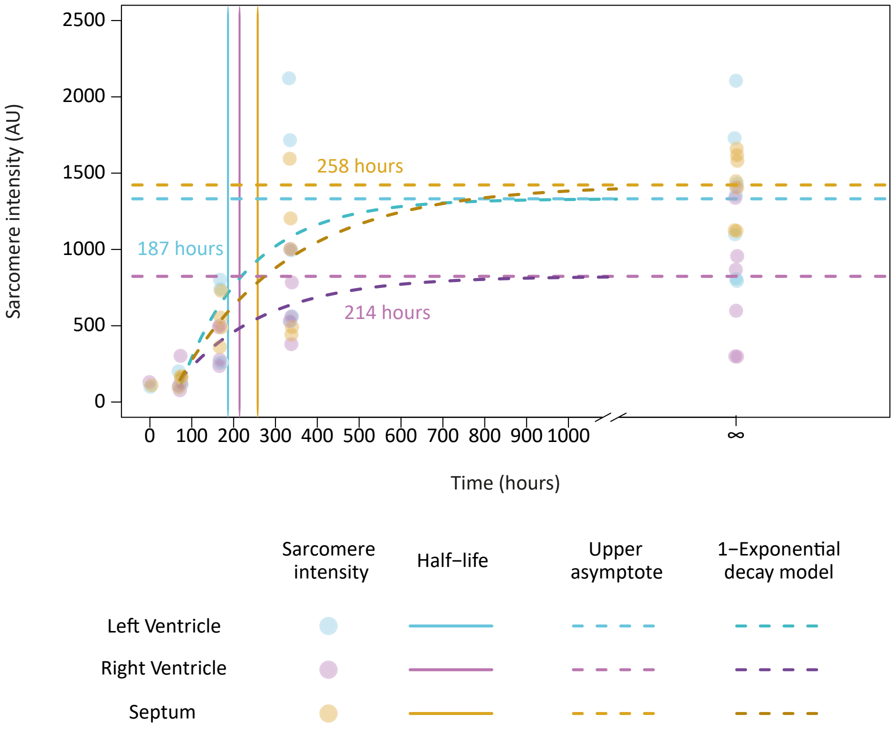
Sarcomere intensity (AU)
2500
2000
1500
1000
500
0
0
100
200
300
400
500
600
700
800
900
1000
∞
Time (hours)
258 hours
187 hours
214 hours
Sarcomere intensity
Half−life
Upper asymptote
1−Exponential decay model
Left Ventricle
Right Ventricle
Septum
