## Supplementary Methods for "Imaging of existing and newly translated proteins elucidates mechanisms of sarcomere turnover"

Please see the Major Resources Table in the Supplemental Materials

**Tissue culture**

Neonatal rat ventricular myocytes (NRVMs) were isolated using the Worthington Biochemical Neonatal Cardiomyocyte Isolation system (cat#LK003300), according to the manufacturer’s instructions, followed by Percoll gradient centrifugation, as previously described.^25^

Neonatal mouse ventricular cardiomyocytes (NMVMs) were isolated according to previously published protocol.^26^ Briefly, hearts were harvested and washed in 1xPBS supplemented with 20 mM BDM. Hearts were then minced and incubated in isolation medium (20 mM BDM, 0.0125% trypsin in HBSS (without Ca2+, Mg2+) overnight at 4°C. The following day the predigested hearts were transferred to a digestion medium (20 mM BDM, Worthington collagenase (cat#LK003240) in L15 medium) and were incubated at 37°C for 25 min with gentle shaking. Tissue fragments were triturated 15 times, and the cell suspension was strained. Cells were centrifuged at 300xg for 5 min. The supernatant was removed, and cells were resuspended in 10 ml of plating medium (65% DMEM high glucose, 19% M-199, 15% fetal calf serum, 1% penicillin/streptomycin).The translation inhibitor cycloheximide (Sigma C7698) was added to myocytes in a concentration of 50 µg/ml for 8 hours.

**Adenovirus vector production and transduction**

The segments containing sarcomeric proteins or truncated protein coding sequence were amplified from a library of genomic NRVM DNA by PCR and cloned into Gateway cloning system (Thermo Fisher) 'Entry' plasmid. The coding sequence was verified by Sanger sequencing and cloned into pAd/CMV/V5-DEST adenovirus expression plasmid (Thermo Fisher). HEK293 cells were transfected with the adenoviral plasmids as previously described^25^, for the generation and propagation of adenoviral particles. Cultured cardiomyocytes were transduced one day after seeding by incubation with the adenovirus for 4 hours, followed by washing by replacing the culture medium.

The constructs used:

- Homo sapiens full size Actinin alpha 2 (ACTN2) – 894 amino acids, with HaloTag in the C terminus. GenBank: KR712245.1
- Rattus norvegicus Myomesin 1 (Myom1) 507 last amino acids, with HaloTag in the C terminus. GenBank: NM_001191584.1
- Rattus norvegicus full size Myosin light chain 2 (Mlc2) – 166 amino acids, with HaloTag in the N terminus. GenBank: NM_001035252.2.
- Rattus norvegicus Myosin binding protein C3 (Mybpc3) 416 last amino acids with HaloTag in the N terminus. GenBank: NM_001106490.1.

**Pulse-chase in cardiomyocyte cultures**

NRVMs or NMVMs were cultured on µ-Dish 35-mm Quad (Ibidi, 80416) coated with Matrigel Matrix (Corning, 354234), in 15% serum media. Cells were transduced with adenoviral vectors encoding the Halo tagged sarcomeric proteins. After 48 hours, 1 µM Oregon Green HaloTag Ligand (Promega G2801) was added to the cells for 15 min. The cells were then incubated with 65µM of 7-bromoheptanol (7BRO, Sigma-Aldrich 310913) in medium for additional 15 min, and washed by replacing the culture medium 3 times. The Janelia Fluor 635 ligand, 0.2 µM, (a kind gift from Luke Lavis) was then added to the cells.

The MG-132 experiments were performed with adenovirus mediated ACTN2-Halo transduced NRVMs, or with NMVM isolated from Halo-TTN knock-in mice. We added 5 µM MG-132 (Apexbio, A2585) or vehicle and imaged over 72 hours. The calpain inhibitor ALLN experiments were performed with NMVM isolated from Halo-TTN knock-in mice. We added 20 µM ALLN (Sigma A6185) or vehicle and imaged over 72 hours.

**Protein degradation experiment**

For degradation experiments NRVM were transduced with adenoviral vector encoding for MYOM1-Halo. After 48 hours cells were incubated with 1 µM Oregon Green HaloTag Ligand for 15 min followed by 65µM 7BRO for 15 min and three washes by replacing the culture medium. Janelia Fluor 635 ligand, 0.2 µM, was added, and cells were incubated for 48 hours to allow synthesis of new proteins and binding of the red ligand. The cells were incubated again with 65µM 7BRO medium, washed 3 times by medium exchange, and imaged over 72 hours.

Alternatively, after cells were transduced and labeled with 1 µM Oregon Green Halotag ligand, the cells were incubated with 65µM 7BRO for 12 hours, followed by three washing by media exchange and labeling with Janelia Fluor 635 ligand 0.2 µM for 24 hours. Then, the cells were incubated again with 65µM 7BRO for 15 minutes, washed three times by medium exchange, and imaged over 72 hours.

**Homology-independent targeted integration**

Neonatal cardiomyocytes primary cultures were prepared from Rosa26-Cas9 knockin mice on C57BL/6J background (Jackson laboratory Strain #:026179).^27^ Cardiomyocytes were transduced with two AAV9 vectors (1 X 10^11^ viral genomes each), one encoding the HaloTag donor cassette and another one encoding for either control non-targeting sgRNA (GATGTTCAAGATTCAGGAGG), or the *Actn2* c-terminus sgRNA (CAGTACTGGGCCTGATCCGG), or the *Myom1* c-terminus sgRNA (TCATCCCAGAGGAGGAGTTG) under the control of the U6 promoter.

**Live cell imaging**

Cells were imaged using Zeiss LSM 900 with Airyscan2 super-resolution system, attached to Axio Observer 7 inverted microscope, with PECON incubation insert, allowing regulation of temperature, CO_2_, and humidity. The acquisition was done with Plan-Apochromat 63x/1.4 Oil DIC M27 (#420782-9900-799) Zeiss objective, with solid state laser with wavelengths of 488 & 640 nm. The photobleach was performed with the 488 nm laser in a small square area at the center of the field.

**Image analysis**

Time series images were aligned using the SIFT-algorithm, in ImageJ software.^28^ For each cell linear regions of interest (ROIs) of ~10 µm were marked, inside and outside the photobleached area, and were used for all the time-lapse images of that cell. In the degradation and in vivo experiments, in which no photobleach was performed, only one ROI was marked and analyzed. The intensity values of the ROI were analyzed using R (R: A language and environment for statistical computing (R Foundation for Statistical Computing, Vienna, Austria. URL [https://www.R-project.org/](https://www.r-project.org/)). We determined the local maxima and minima points along the ROI using a moving window of 20 pixels (~1 µM). The amplitudes along the ROI were calculated from the local maxima and minima, and the median of all amplitude values was used as the representative value for the myofibril or cell. To calculate the half-life the median amplitude values from each cell were normalized between 0 and 1 to standardize the values and allow analysis of all cells together. Nonlinear Least Squares (NLS) function was used to fit all the representative values according to equation 1 for protein degradation, in exponential decay model, and equation 2 for protein synthesis, in 1 minus exponential decay model^29^.

$$Eq. 1: intensity=K*e^{(-t*\lambda)}$$

$$Eq.2: intensity=1-K*e^{(-t*\lambda)}$$

K and $\lambda$ are parameters estimated by the NLS function, *t* is the time and *intensity* is the calculated representative value.

The half-life calculated for each fitting model was calculated according to equation 3.

$$Eq.3: t½= \frac{\ln(2)}{\lambda}$$

The R^2^ values for the fitting to exponential curves were calculated by R for the linear model log(y)~ x

**Adeno-associated virus vector production and injection**

The AAVs were produced according to a published protocol^30^, as we previously described.^31^ For transduction 50µl of virus containing a total of ~3 X 10^11 viral genomes was delivered intraperitoneally to 4-7-days old mouse pups.

Mouse pups were injected with AAV viruses, encoding for the following constructs or control (no virus):

- Homo sapiens full size Actinin alpha 2 (ACTN2) – 894 amino acids, with HaloTag in the C terminus. GenBank: KR712245.1
- Rattus norvegicus full size Tropomyosin 1 (TPM1) - 285 amino acids, with HaloTag in the C terminus. GenBank: NM_001301336.1

**Animal experiments**

All pups were born at the SPF unit of the pre-clinical research authority at the Technion. Health monitoring was carried out in accordance with FELASA recommendations. Each litter was housed with the dam weened at 4 weeks and separated by sex. Mice were maintained under climate-controlled conditions of 12:12 – hours light/dark cycle temperature range 21±2^o^C, relative humidity of 30-70% and fed ad libitum a commercial food pellet diet. Studies were conducted at the Technion, (IIT), Faculty of Medicine, Haifa, Israel, after obtaining approval from the institute’s IACUC. All proceedings complied with the Animal Welfare Act of 1966 (P.L. 89-544), as amended by the Animal Welfare Act of 1970 (P.L.91-579) and 1976 (P.L. 94-279).

All results are reported in compliance with the Animal Research Reporting of In Vivo Experiments (ARRIVE) guidelines. Mice were randomly assigned to each experimental group. Male and female mice were equally used. No mice were excluded.

7-bromoheptanol (Sigma-Aldrich, 310913) (7BRO) 325 mM in DMSO (Sigma-Aldrich, D2650) was injected to adult mice retro-orbitally (15 µl) and intraperitoneally (50 µl) to achieve complete blocking. (Sigma-Aldrich, D2650), or intraperitoneally only to neonatal mice.

Janelia Fluor-525 HaloTag ligand was a kind gift from Luke D. Lavis. One µl of 1µM stock solution was diluted in 15µl DMSO and injected retro-orbitally. The retro-orbital injections were done under sedation with isoflurane (Piramal, Isoflurane, USP Terrell).

**In-vivo AAV overexpression**

Adult mice (4 month old) received 7BRO (or control no 7BRO). After either 3 hours, 1, 3, 7 days the mice received Janelia Fluor-525 HaloTag ligand retro-orbitally. Mice were sacrificed, and hearts were harvested 40 minutes after the ligand injection.

**In-vivo experiments in HaloTag-TTN knock-in mice**

The HaloTag-TTN knock-in mice were previously described.^32^ Adult mice (4-month-old) received 7BRO (or control no 7BRO). After either 3 hours, 3, 7, or 14 days the mice received the Janelia Fluor-525 HaloTag ligand. Mice were sacrificed, and hearts were harvested 40 minutes after the ligand delivery. For hypertrophy, 9mM PE (Sigma-Aldrich, P6126) and 0.2% ascorbic-acid (Sigma-Aldrich, 95209) in PBS were filtered through a 0.22 µm filter. Mice received three subcutaneous injections of 10 mg/kg bodyweight PE on days -7, -5, and -3, with day 0 being the day of the sacrifice. For experiments in neonatal mice 1-day-old pups received 7BRO (or control no 7BRO). After either 3 hours, 1, 3, 7, or 10 days the mice were sacrificed, and hearts were harvested.

**Heart sections preparation**

After harvest, hearts were incubated in 4% formaldehyde for 1-2 hours on ice, and then moved to overnight incubation with gentle tilting in 50% sucrose in PBS solution. Next the hearts were submerged in Tissue Freezing Medium (Leica, 7876270), and frozen in isopentane cooled in liquid nitrogen. Cryosection was carried out by Leica CM1860 UV Cryostat, 5µm sections were performed, and mounted on a slide. Sections were then fixed with 4% formaldehyde for 10 min at room temperature, permeabilized with 1% Triton X-100 (Sigma-Aldrich) in PBS for 10 min, nuclear counterstained with DAPI (Sigma-Aldrich, D9542) at 1µg/ml in PBS, mounted with 15µl of Fluoromount G (Enco, 0100-01) and sealed with optic round cover slides. For the TTN knock-in mice sections were washed twice with PBS containing 0.3% Triton X-100 (Sigma-Aldrich), and incubated with Janelia Fluor-525 HaloTag ligand for 30 minutes at 37^o^ Celsius. Then, sections were washed once again with PBS containing 0.3% Triton X-100 (Sigma-Aldrich), nuclear counterstained with DAPI (Sigma-Aldrich, D9542) at 1µg/ml in PBS, mounted with 15µl of Fluoromount G (Enco, 0100-01) and sealed with optic round cover slides.

**Heart sections imaging**

Heart sections were imaged using Zeiss LSM 880 with Airyscan super-resolution system, attached to Axio Examiner Z1 microscope. The acquisition was done with Plan-Apochromat 63x/1.4 Oil DIC M27 (#420782-9900-799) Zeiss objective, with diode laser 405nm wavelength and argon laser 514 nm wavelength. Images were then Airyscan processed with ZEN software (Zeiss) using the standard parameters.

**In-vivo image analysis**

A linear ROI of several sarcomeres' length was marked on every cell, and the signal intensity values were extracted. The representative value was calculated the same way as in the live cells imaging analysis. We did not include the 0- and 3-hours controls (background) data in the fitting because of the low signal to noise ratio at the early time points. We fitted the data with a 1-exponential decay model, limited to a maximal value of the average of the No 7BRO control group (*c*), as this control represents the steady-state protein level (Eq.4)

$$Eq.4: intensity=c-K*e^{(-t*\lambda)}$$

**Western Blot analysis**

Proteins were resolved by SDS-PAGE, probed with either anti-Actn2 antibody (Sigma, A7811) or anti-Ubiquitin antibody (Cytoskeleton, AUB01), and visualized using the LI-COR Biosciences Odyssey M system. Quantification of bands was done using ImageJ.

**Detection of HaloTag fusion proteins using SDS-PAGE**

NRVMs or NMVMs were cultured on 6-well plates (Thermo Scientific, 140675) coated with Matrigel Matrix (Corning, 354234), in 15% serum media. Cells were transduced with adenoviral vectors encoding the Halo tagged sarcomeric proteins. After 48 hours, 1 µM Oregon Green HaloTag Ligand (Promega G2801) was added to the cells for 15 min. The cells were then incubated with 65µM of 7-bromoheptanol (7BRO, Sigma-Aldrich 310913) in medium for additional 15 min, and washed by replacing the culture medium 3 times. After either 0, 24, 48 hours 0.1 µM TMRDirect HaloTag Ligand (Promega G2991) was added to the cells. Cells were then washed with 1X PBS (pH 7.5). Cells were lysed by replacing 1X PBS with 150ul of 1X SDS sample buffer. Lysate was incubated for 5 minutes at 95°C, and resolved by SDS-PAGE. Gels were analyzed on the LI-COR Biosciences Odyssey M system. Quantification of bands was done using ImageJ.

**Global fitting analysis**

Data was analyzed using an independent fit method or a global fit method using Origin (Version 2024, OriginLab Corporation, USA).

**Statistics**

Data are shown as mean ± SE as indicated. A two tailed student's t-test was used for comparison, unless otherwise specified.
